# Prolyl tRNA Synthetase Is Required for Mammarenavirus Multiplication

**DOI:** 10.64898/2026.01.15.699610

**Authors:** Haydar Witwit, Pablo Ibanez, Ruifeng Zhou, Nathaniel Jackson, Ruby Escobedo, Beatrice Cubitt, Roaa Khafaji, Rachel Y. Sattler, Luis Martinez-Sobrido, Juan Carlos de la Torre

## Abstract

Several mammarenaviruses (MaAv), chiefly Lassa virus (LASV) in Western Africa and Junin virus (JUNV) in the Argentinean Pampas, cause severe disease in humans and pose important public health problems in their endemic regions. In addition, the globally distributed MaAv lymphocytic choriomeningitis virus (LCMV) is an underrecognized human pathogen of clinical significance especially in congenital infections and LCMV poses a serious risk for immunocompromised individuals. There are no FDA-approved MaAv vaccines or antivirals and current anti-MaAv therapy is limited to an off-label use of ribavirin whose efficacy remains controversial. This highlights an urgent unmet need for developing antivirals against human pathogenic MaAv. Halofuginone (HF), a derivative of the natural alkaloid febrifugine, has been shown to exhibit antiviral activity against several RNA viruses. Here, we present evidence that HF exhibits a potent dose-dependent antiviral activity against LCMV, and the hemorrhagic fever causing MaAv LASV and JUNV. HF binds to the bifunctional enzyme glutamyl-prolyl-tRNA synthetase 1 (EPRS1) and specifically inhibits its prolyl-tRNA synthetase (PRS) activity, resulting in translation inhibition via the amino acid starvation (AAS) response with preferential impact on proline-rich proteins. HF anti-LCMV activity was prevented by the addition of exogenous proline supporting that inhibition of PRS activity plays a critical role on the anti-MaAv activity of HF. We found that HF did not affect LCMV cell entry, modestly (twofold) reduced the activity of the virus ribonucleoprotein (vRNP) but strongly inhibited (>90%) Z budding activity, a process involving the Z proline-rich late domain motifs.

## 1. Introduction

Mammarenaviruses (MaAv) cause persistent infections in multiple rodent species worldwide, and human infections occur primarily through mucosal exposure to aerosols or direct contact of abraded skin with infectious materials. Several MaAv cause viral hemorrhagic fever disease (VHFD) in humans and represent important public health challenges in their endemic regions. Thus, the MaAv, Lassa virus (LASV) is highly prevalent in Wester Africa where it is estimated to infect over 500,000 people annually [1], resulting in a high number of Lassa fever (LF) cases, a VHFD associated with high morbidity, and case fatality rates as high as 69% among hospitalized confirmed LF cases [2–4]. LASV endemic regions are expanding [5], and increased global travel has led to the importation of LF cases into non-endemic metropolitan areas [6,7]. Notably, LASV ranks very high among zoonotic viruses with potential for spillover and spread in humans [8]. Likewise, the MaAv Junin virus (JUNV) causes Argentine hemorrhagic fever with case fatality rates exceeding 15%, and several other New World MaAv cause VHFD throughout South America [9,10]. Moreover, several MaAv, including LASV and JUNV, are credible biodefense threats and are classified Category A agents [11]. In addition, mounting evidence indicates that the worldwide-distributed MaAv lymphocytic choriomeningitis virus (LCMV) is a neglected human pathogen of clinical significance in congenital infections and transplantation medicine [12–14]. Although most acquired LCMV infections in immunocompetent are asymptomatic or result in self-limited mild illness [15], in some cases LCMV infected immunocompetent individuals can develop severe disease [16].

There are currently no FDA-approved MaAv vaccines or antivirals, and current anti-MaAv therapeutic options are limited to an off-label use of ribavirin whose efficacy remains controversial [17]. A monoclonal antibody cocktail has shown protection in preclinical models of LF [18], but high cost, cold chain distribution and storage requirements, and parenteral delivery, may limit their use to specialized treatment centers. The broad-spectrum polymerase inhibitor nucleoside analogs favipiravir [19–22], and 4’-fluorouridine (4’-FIU) [23,24], as well as cell entry inhibitors LHF-535 [25], and ARN-75039 [26] that target the fusion activity of the virus surface glycoprotein (GP) have shown promising results in preclinical models of LF, and LHF-535 is in phase I clinical trials. However, favipiravir did not show superior efficacy over ribavirin in a phase II clinical trial for the treatment of LF [27], whereas the linkage of 4’-FIU to mitochondrial dysfunction and cardiotoxicity may complicate its clinical development. Likewise, the high genetic diversity of LASV [28] can compromise cell entry inhibitors as single mutations in GP can confer cross-resistance to these type of antivirals [29]. Thus, the development of broad-spectrum antivirals against human pathogenic MaAv represents a significant unmet biomedical need.

Halofuginone (HF), an FDA-approved feed additive derived from the natural alkaloid febrifugine, is a potent competitive inhibitor of the prolyl tRNA synthetase (PRS) activity of the bifunctional enzyme glutamyl-prolyl-tRNA synthetase 1 (EPRS1) that is currently under investigation as a therapeutic for the treatment of fibrotic disorders, cancer and autoimmune diseases [30–32]. Notably, HF has also been shown to exhibit potent antiviral activity against different viruses, including CHIKV, DENV, and SARS-CoV-2 [33,34]. Here we present evidence that HF exhibits a potent antiviral activity against LCMV and the hemorrhagic fever causing MaAv LASV and JUNV. HF mediated inhibition of PRS results in inhibition of translation via the amino acid starvation (AAS) response with a higher impact on proline-rich proteins [35]. HF anti-LCMV activity was prevented by the addition of exogenous proline supporting that inhibition of PRS activity plays a critical role on the anti-MaAv activity of HF. MaAv matrix Z protein is the main driving force of viral budding, a process involving the Z proline-rich late domain motifs, which may make Z budding activity highly vulnerable to HF. Consistent with this hypothesis, we found that HF did not affect LCMV cell entry but rather modestly (two-fold) reduced the activity of the virus ribonucleoprotein (vRNP), responsible for directing replication and transcription of the viral genome, but drastically inhibited (>90%) Z budding activity.

## 2. Materials and Methods

### 2.1. Cells and viruses

African green monkey kidney, Vero E6, cells (ATCC CRL-1586), and human A549 (ATCC CCL-185), HEK 293T (ATCC CRL-3216), and Ea HY 926 (ATCC CRL-2922) cell lines were cultured in Dulbecco’s Modified Eagle Medium (DMEM; ThermoFisher Scientific, Waltham, MA, USA) supplemented with 10% heat-inactivated fetal bovine serum (FBS), 2 mM L-glutamine, 100 µg/mL streptomycin, and 100 U/mL penicillin. HAP1 cells (Horizon Discovery) were maintained in Iscove’s Modified Dulbecco’s Medium (IMDM) supplemented with 10% FBS and 100 µg/mL streptomycin, and 100 U/mL penicillin. Recombinant viruses: rLCMV/GFP-P2A-NP (referred to as rLCMV/GFP), expressing green fluorescent protein (GFP) fused to the LCMV nucleoprotein (NP) via a P2A ribosomal skipping sequence; rLCMVΔGPC/ZsG-P2A-NP (rLCMVΔGPC/ZsG)[36], a single-cycle variant expressing Zoanthus sp. ZsG; the tri-segmented JUNV Candid#1 strain expressing GFP (r3Can/GFP) [37]; a tri-segmented form of the LASV Josiah strain expressing GFP (r3LASV/GFP) [38]; and rLCMV/Z-HA [39] have been described. All the experiments with LCMV were conducted in the biosafety level 2 (BSL-2) laboratory at The Scripps Research Institute. All the experiments with r3LASV and r3JUNV were conducted in the biosafety level 4 (BSL-4) laboratory at Texas Biomedical Research Institute.

### 2.2. Antibodies and compounds

Halofuginone hydrobromide (Cat. No. HY-N1584A; MedChemExpress, NJ, USA) was dissolved in DMSO at 10 mM and stored in aliquots at −20 °C. F3406-2010 was obtained from DC Chemicals (Cat. No. DC11877). Ribavirin (Cat. No. R9644) and L-proline (Cat. No. 81709-10G) were purchased from Sigma-Aldrich (St. Louis, MO, USA). IMP-1088 (Cat No. 25366-1) was purchased from Cayman Chemical, dissolved in methyl acetate at 11 mM, and kept in aliquots at −20. Sodium citrate buffer (0.1 M, pH 5.0, sterile) was obtained from bioWorld (MFG No. 40121003-1, Dublin, OH, USA). Anti-HA antibody was purchased from GenScript (Piscataway, NJ, USA); the VL4 rat monoclonal antibody against NP was obtained from Bio X Cell (West Lebanon, NH, USA).

### 2.3. Cell viability (CC₅₀) and viral inhibition half maximal effective concentration (EC₅₀)

Cell viability was assessed using the CellTiter 96 AQueous One Solution Reagent (Promega, Madison, WI, USA), which quantifies viable cells based on the conversion of a tetrazolium compound to a formazan product by NADPH or NADH in metabolically active cells. A549 cells were seeded in 96-well clear-bottom plates (4.0 × 10⁴ cells/well), treated with 2-fold serial dilutions of each compound, and incubated for 72 h. Following treatment, CellTiter reagent was added and incubated for 35 min at 37 °C in 5% CO₂. Absorbance was measured using a Cytation 5 reader (BioTek, Winooski, VT, USA), and values were normalized to vehicle (DMSO)-treated controls, set at 100%. The half-maximal cytotoxic concentration (CC₅₀) was calculated using GraphPad Prism v10 (Prism10). Cell viability was also determined using DAPI staining. For determination of the half-maximal effective concentration (EC₅₀), A549 cells were seeded in 96-well clear-bottom black plates (4.0 × 10⁴ cells/well), and 20 h later infected (MOI = 0.05) with rLCMV/GFP-P2A-NP. After 90 min of adsorption, the viral inoculum was removed and replaced with media containing test compounds. At 72 h post-infection, cells were fixed with 4% paraformaldehyde (PFA), and GFP expression was measured by fluorescence using a Cytation 5 reader. Fluorescence readings were normalized to DMSO-treated controls (100%), and EC₅₀ values were calculated using Prism10. The selectivity index (SI) for each compound was determined as the ratio of CC₅₀ to EC₅₀.

### 2.4. Viral growth kinetics

Cells were infected in 12-well plates at the indicated cell seeding density and MOI. After 90 min of adsorption at 37 °C and 5% CO₂, the virus inoculum was removed, cells were washed once with DMEM containing 2% FBS, and fresh media with the indicated compounds and concentrations was added. At specified times post-infection cell culture supernatants (CCS) were collected, and viral titers were determined by focus-forming assay (FFA).

### 2.5. Virus titration

Virus titers were determined by focus-forming assay (FFA) as described [40]. Briefly, serial 10-fold dilutions of samples were prepared in DMEM containing 2% FBS and used to infect Vero E6 cell monolayers seeded in 96-well plates (2 × 10⁴ cells/well). At 20 h post-infection, cells were fixed with 4% paraformaldehyde in phosphate-buffered saline (PBS), and infection foci were visualized by epifluorescence based on GFP expression from rLCMV/GFP-infected cells using a fluorescence microscope.

### 2.6. LCMV and LASV cell-based minigenome (MG) assay

The LCMV cell-based MG assay was performed as described [41], but transfecting the cells in suspension. HEK293T single cell suspension were prepared to obtain a total of 4.84 × 10⁶ cells in 5.2 mL of complete medium in a 50 mL conical tube. A transfection mixture (TM; 800 µL) was added to reach a final volume of 6 mL, and the suspension was incubated for 10 min at room temperature. Following incubation, 3.6 mL of complete medium was added and mixed gently by pipetting to ensure uniform distribution, resulting in a final TM volume of 9.6 mL. TM was added (80 µL/well) to poly-L-lysine coated wells (96 well-plate) pre-spotted with 20 µL of 5x HF. Wells were mixed twice with an electronic pipette, and plates were placed on a level surface platform for 15 min before being transferred to an incubator at 37°C in a humidified atmosphere with 5% CO₂.

The TM consisted of Lipofectamine 3000 (2.5 µL/µg DNA; ThermoFisher Scientific) with Pol II-driven expression plasmids encoding T7 RNA polymerase (pC-T7, 1.32 µg), nucleoprotein (NP; pC-NP, 0.19 µg), and L polymerase (pC-L, 1.58 µg), along with a T7 promoter-driven plasmid expressing an LCMV S MG encoding the GFP reporter (pT7-MG/GFP, 1.32 µg) (Fig. 1). The LASV MG transfection mixture was prepared by combining 403.75 µL of Opti-MEM containing plasmid DNA and 25 µL of P3000 reagent with 403.75 µL of Opti-MEM containing 37.5 µL of Lipofectamine 3000, yielding a total volume of 806.6 µL. The mixture was incubated for 20 min at room temperature to allow for complex formation. At the indicate end timepoint, the cells were fixed with 4% paraformaldehyde (PFA), washed with PBS, then stained with DAPI. Images and GFP signal readings were collected using Keyence BZ-X710 and Cytation 5 reader respectively.

**Figure 1.**
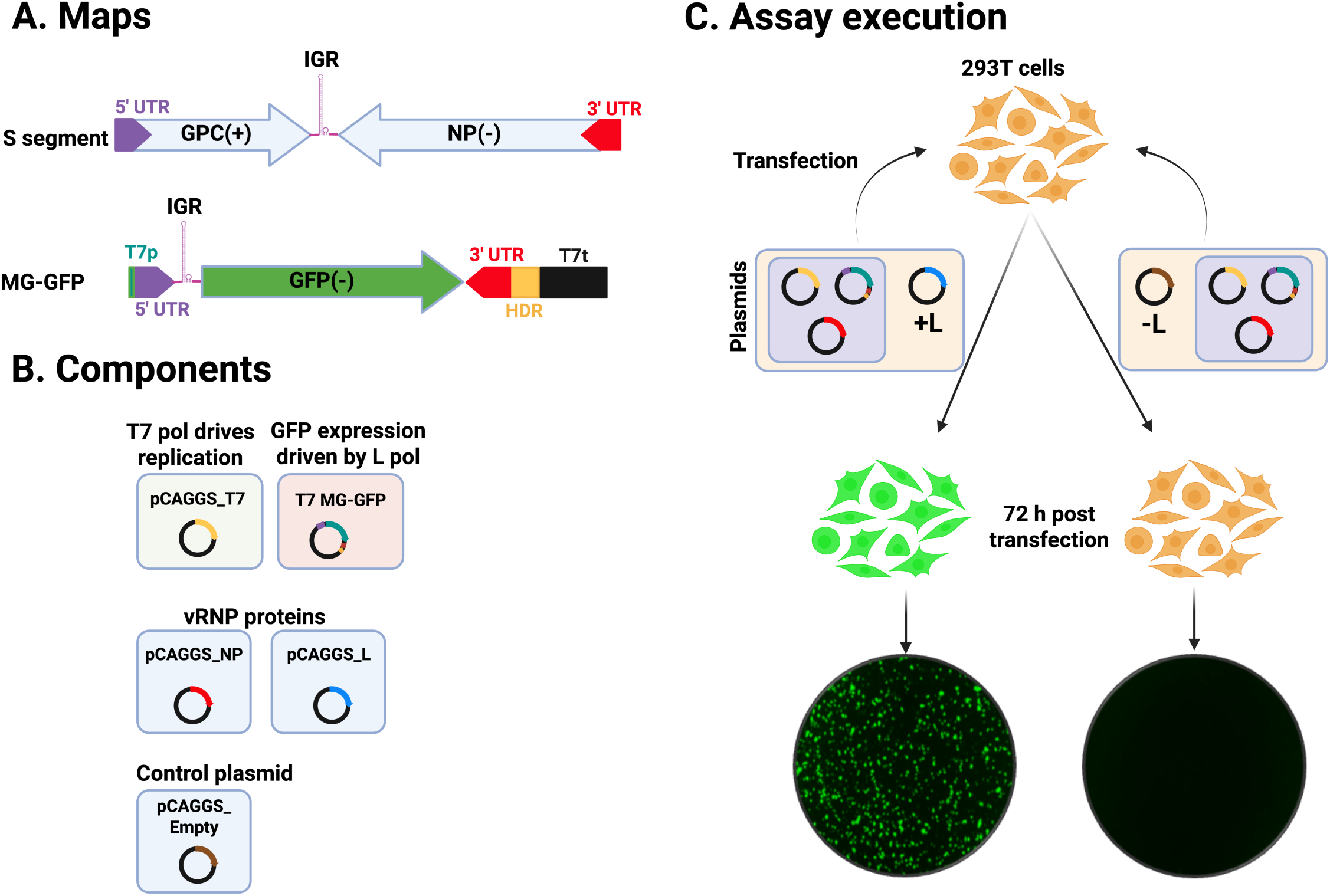
Schematic of the components and steps of the cell-based minigenome assay. **A.** Schematic of the LCMV S genome segment and the S-based minigenome (MG) expressing GFP from the NP locus. **B.** Components of the LCMV cell-based MG system based on the use of T7 RNA polymerase to initially launch intracellular synthesis of the MG RNA. **C.** Schematic of the transfection protocol use in the cell-based MG assay and detection of MG-directed expression of GFP. Green animations and IF images indicate GFP expression, whereas orange animations and black IF images indicate no expression. T7, T7 polymerase; L, L polymerase; NP, nucleoprotein; GPC, glycoprotein precursor; UTR, untranslated region; IGR, intergenic region; HDR, hepatitis D ribozyme; T7p, T7 promoter; T7t, T7 terminator; (−), negative polarity; (+), positive polarity. Images and animations are not to scale.

### 2.7. Time of addition assay

Vero E6 cells were seeded in 96-well plates at a density of 2 × 10⁴ cells/well. The following day, cells were infected with the single-cycle infectious rLCMVΔGPC/ZsG (MOI = 0.5) and treated with HF (75 nM) or vehicle control (VC) either 2 h before infection (−2 h) or 2 h post-infection (+2 h). The LCMV entry inhibitor F3406 (10 µM) was included as a control treatment. At 48 h pi, ZsGreen-positive (ZsG⁺) cells were quantified using the Cytation 5 Reader (BioTek, Agilent). Fluorescence values were normalized to those of VC-treated infected cells, and results represent the mean of three biological replicates.

### 2.8. GPC-mediated fusion assay

HEK293T cells were seeded in poly-L-lysine-coated 24-well plates at a density of 2.5 × 10⁵ cells/well. The following day, cells were transfected with a pCAGGS plasmid encoding GFP (50 ng/well) along with either an empty pCAGGS vector or pCAGGS plasmids expressing LCMV or LASV GPC (1 µg/well) using Lipofectamine 3000, following the manufacturer’s instructions. After 5 h, the transfection medium was removed, cells were washed once, and fresh medium with or without HF was added. At 24 h post-transfection, cells were exposed to either acidified (pH 5.0) or neutral (pH 7.2) medium for 15 min, washed with DMEM, and returned to DMEM containing 10% FBS. Cells were then monitored over time for syncytium formation using fluorescence microscopy. Following observation, cells were fixed with 4% PFA, washed with PBS, and imaged at 20X magnification using a Keyence BZ-X710 microscope.

### 2.9. Z budding assay

Z budding activity was assessed using a described cell-based Z budding assay [38]. HEK293T cells were seeded in poly-L-lysine-coated 12-well plates at a density of 3.5 × 10⁵ cells/well. After overnight incubation, cells were transfected with 2 µg of either pC-LCMV-Z-Gaussia luciferase (GLuc), pC-LCMV-mutant Z [G2A]-GLuc, or pC-LASV-Z using Lipofectamine 2000 (2.5 µL/µg DNA). Following a 5 h incubation, the transfection mixtures were replaced with fresh media containing the indicated compounds. At 48 h post-transfection, CCSs containing virion-like particles (VLPs) were collected and clarified by low-speed centrifugation to remove cell debris. Aliquots (20 µL) of each CCS sample were transferred to 96-well black plates (VWR, West Chester, PA, USA), followed by the addition of 50 µL Steady-Glo luciferase reagent (Promega). Whole-cell lysates (WCLs) from the same wells were prepared to assess cell-associated GLuc activity. Luminescence was measured using the Cytation 5 Reader (BioTek, Agilent). Budding efficiency was calculated as CCS-GLuc/CCS-GLuc + WCL-GLuc.

### 2.10. Assessment of NP:Z ratio

A549 cells were seeded in 96-well clear-bottom plates at a density of 4.0 × 10⁴ cells/well, infected (MOI = 1) with rLCMV/Z-HA and treated with 3-fold serial dilutions of the indicated compound. At 72 h pi, cells were fixed with 4% PFA and examined by immunofluorescence (IF) using the VL4 rat monoclonal antibody against NP (Bio X Cell, West Lebanon, NH, USA) conjugated to Alexa Fluor 488, and an anti-HA antibody (Cat. No. A01244, GenScript, Piscataway, NJ, USA) conjugated to Alexa Fluor 647 for Z detection. Nuclei were visualized by DAPI staining. Fluorescence signals were normalized to the vehicle-treated (DMSO) control group, set at 100%, and data were plotted using GraphPad Prism v10.

### 2.11. LASV and JUNV Cell viability (CC50) and viral inhibition half maximal effective concentration (EC50)

A549 cells (96-well plate format, quadruplicates) were seeded at 2 x 10^4^ / well, next day, cells were infected with r3LASV (MOI 0.003) or r3JUNV (MOI 0.006). After 1 h of viral adsorption (100µL), indicated concentrations of inhibitors (2x in 100µL) were added (total 200µL of 1x final concentration). At 72 h (r3LASV) or 96 h (r3JUNV) post-infection, cell culture supernatants (CCSs) were collected and cells fixed with 10% formalin. Gluc activity in CCSs was quantified according to the manufacturer protocol using Pierce™ Gaussia Luciferase Glow Assay Kit (16160, Thermo Fisher Scientific) and GloMax plate reader (Promega). Data were normalized to infected and vehicle control treated cells. Fixed cells were imaged for GFP and DAPI expression using an EVOS M5000 imaging system (Thermo Fisher Scientific). DAPI expression was quantified by Syngergy H1 plate reader (Agilent) and was normalized to mock-infected cells to determine cell viability.

## 3. Results

### 3.1. Dose-dependent effect of HF on LCMV multiplication

HF exhibited a potent dose-dependent inhibitory effect on rLCMV/GFP-P2A-NP multiplication in A549 cells, with EC50 and EC90 values of 16 and 26 nM, respectively, at 48 h pi, and 21 and 37 nM at 72 h pi (Fig. 2). HF exhibited a similar anti-LCMV activity in endothelial cells with EC50 and EC90 values of 10 and 21 nM, respectively (SF1). The inhibitory effect of HF on LCMV multiplication was not a consequence of drug-induced cell toxicity, as HF exhibited an CC50 value >10 µM and selectivity index (SI = CC50/EC50) of > 400 (Fig. 2). We obtained similar results in human endothelial Ea.hy926 cells (SF1).

**Figure 2.**
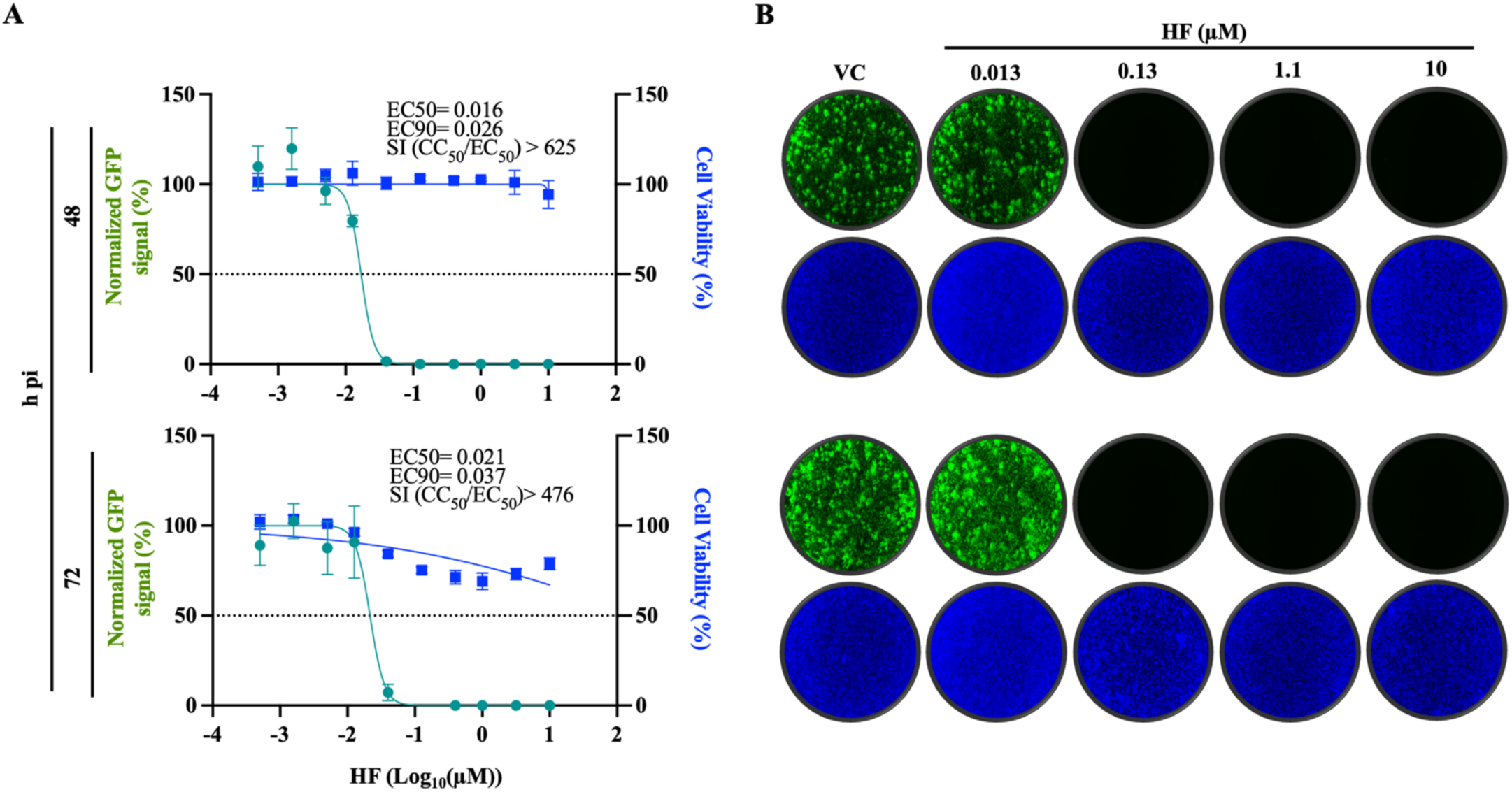
Dose-dependent effect of HF on LCMV multiplication in A549 cells. **A.** A549 cells were seeded at 4 × 10⁴ cells/well into a 96-well plate, infected (MOI = 0.05) with rLCMV/WT-GFP-P2A-NP and treated with HF at the indicated concentrations. At 48 and 72 h pi, cells were fixed with 4% PFA and stained with DAPI to assess cell viability. GFP and DAPI signals were quantified using Cytation 5 normalized (%) to VC-treated infected controls. Results show the mean and SD of four biological replicates. EC50 and CC50 values were calculated using a variable slope (based on four parameters) model and EC90 values were calculated using FindECanything model (logEC50=logECF - (1/HillSlope)*log(F/(10-F)) with F parameter set to 10eq (Prism10). **B.** Representative IF images of the dose-response assay of selected doses of HF. DAPI staining was used to visualize nuclei. Images were taken at 4× magnification using a Keyence BZ-X710 microscope.

### 3.2. Effect of HF on LCMV multi-step growth kinetics

To examine the effect of HF on LCMV multi-step growth kinetics, we infected A549 cells with rLCMV/GFP-P2A-NP (MOI = 0.05) and treated them with HF (100 nM) or vehicle control (VC). At the indicated h pi, we collected CCS samples and determined virus titers using a focus-forming assay (FFA). Treatment with HF abrogated production of infectious viral progeny at all time points tested (Fig. 3A), which correlated with restricted virus propagation within the infected cell monolayer (Fig. 3B, Ci). To assess the effect of HF on cell viability, we quantified DAPI staining as a surrogate of cell number. HF exerted a modest cytostatic effect on A549 cells resulting in ∼10-15% lower cell number at 72 h pi compared to VC treated cells (Fig. 3Cii). Production of infectious viral progeny was also abrogated by treatment with 75 nM HA, whereas low levels (≤ 10^3^ FFU/mL) of infectious viral progeny was detected at 72 h pi in LCMV-infected cells treated with 50 nM HA (Fig. 3D). Results from LCMV multi-step growth kinetics assay were reproduced in independent biological replicates (SF2).

**Figure 3.**
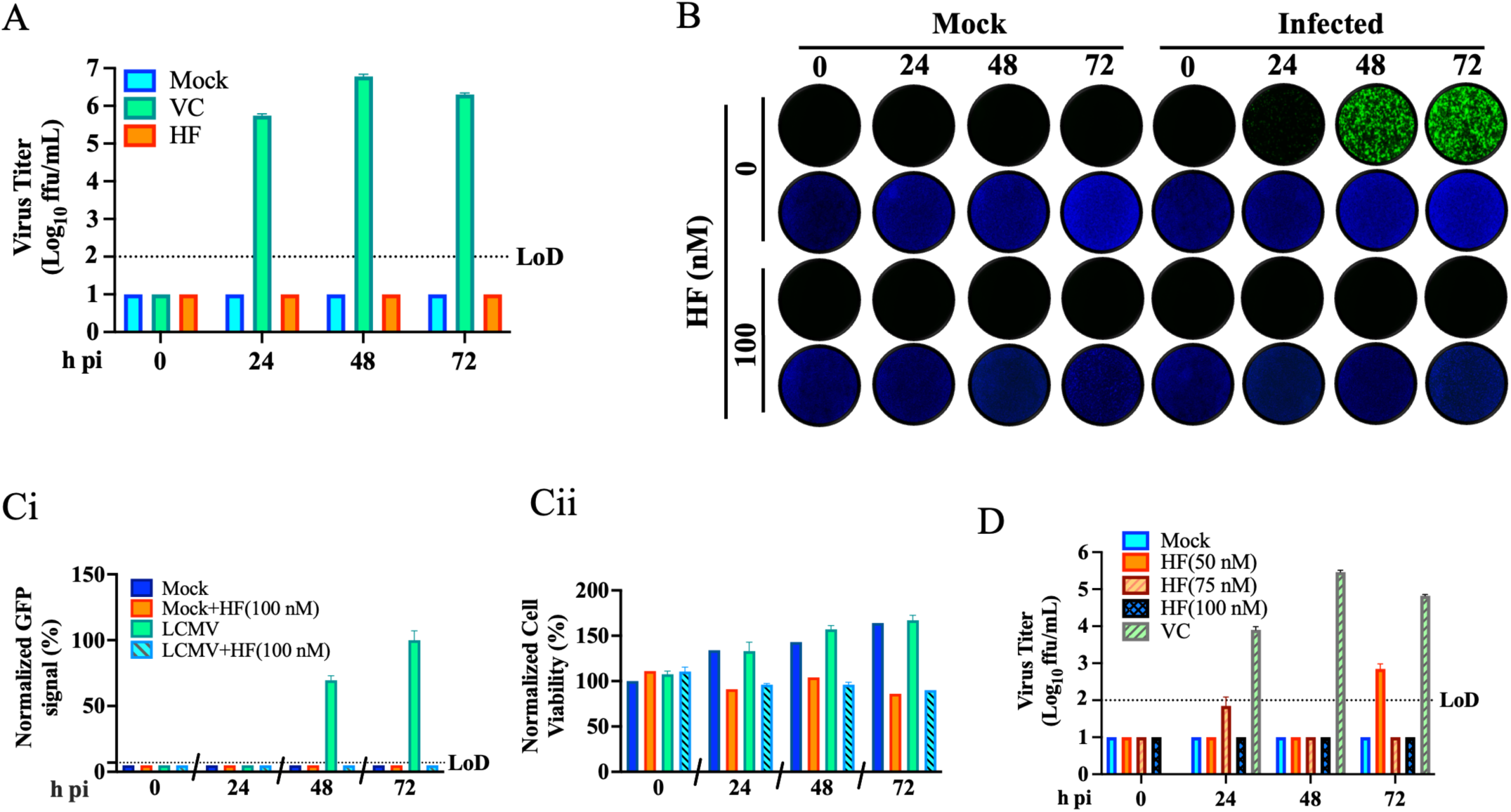
Effect of the HF on LCMV multi-step growth kinetics in A549 cells. **A.** Effect of HF on production of infectious viral progeny. A549 cells were seeded at 5 × 10⁵ cells/well in an M12-well plate, infected (MOI = 0.05) with rLCMV/GFP-P2A-NP and treated with HF (100 nM) or with VC. At the indicated time points, CCSs were collected and titers of infectious virus were determined by FFA using Vero E6 cells. **B.** At the indicated h pi, samples from **A** were fixed with 4% PFA and stained with DAPI. Images (4x magnification) were collected using a Keyence BZ-X710 microscope. **C.** GFP (**Ci**) and DAPI (**Cii**) signals were quantified using Cytation 5 and normalized to the average value of the VC treated samples at 72 h pi. **D.** Dose-dependent effect of HF on LCMV multi-step growth kinetics in A549 cells. A549 cells were seeded at 5 × 10⁵ cells/well in an M12-well plate, infected (MOI = 0.05) with rLCMV/GFP-P2A-NP and treated with HF (50, 75, and 100 nM) or with VC. At the indicated time points, CCS were collected, and titers of infectious virus were determined by FFA using Vero E6 cells.

### 3.3. Effect of proline supplementation on the anti-LCMV activity of HF

To investigate whether the anti-LCMV activity of HF was mainly due to its ability to act as a competitive inhibitor of the PRS activity of EPRS1, we examined the effect of proline supplementation. For this, we infected (MOI = 0.05) A549 cells with rLCMV/GFP-P2A-NP and treated them with HF (75 nM) or VC in the absence or presence of 1 mM L-proline (Pro). At the indicated h pi, we collected CCS samples and determined virus titers using a FFA. Pro treatment reversed the HF inhibitory effect on production of infectious LCMV progeny (Fig.4A), which correlated with the restricted propagation of LCMV within the infected cell monolayer (Fig. 4B, Ci). Consistent with results shown in Fig. 3Cii, treatment with HA in the absence of Pro was associated with a modest cytostatic effect (Fig. 4Cii).

**Figure 4.**
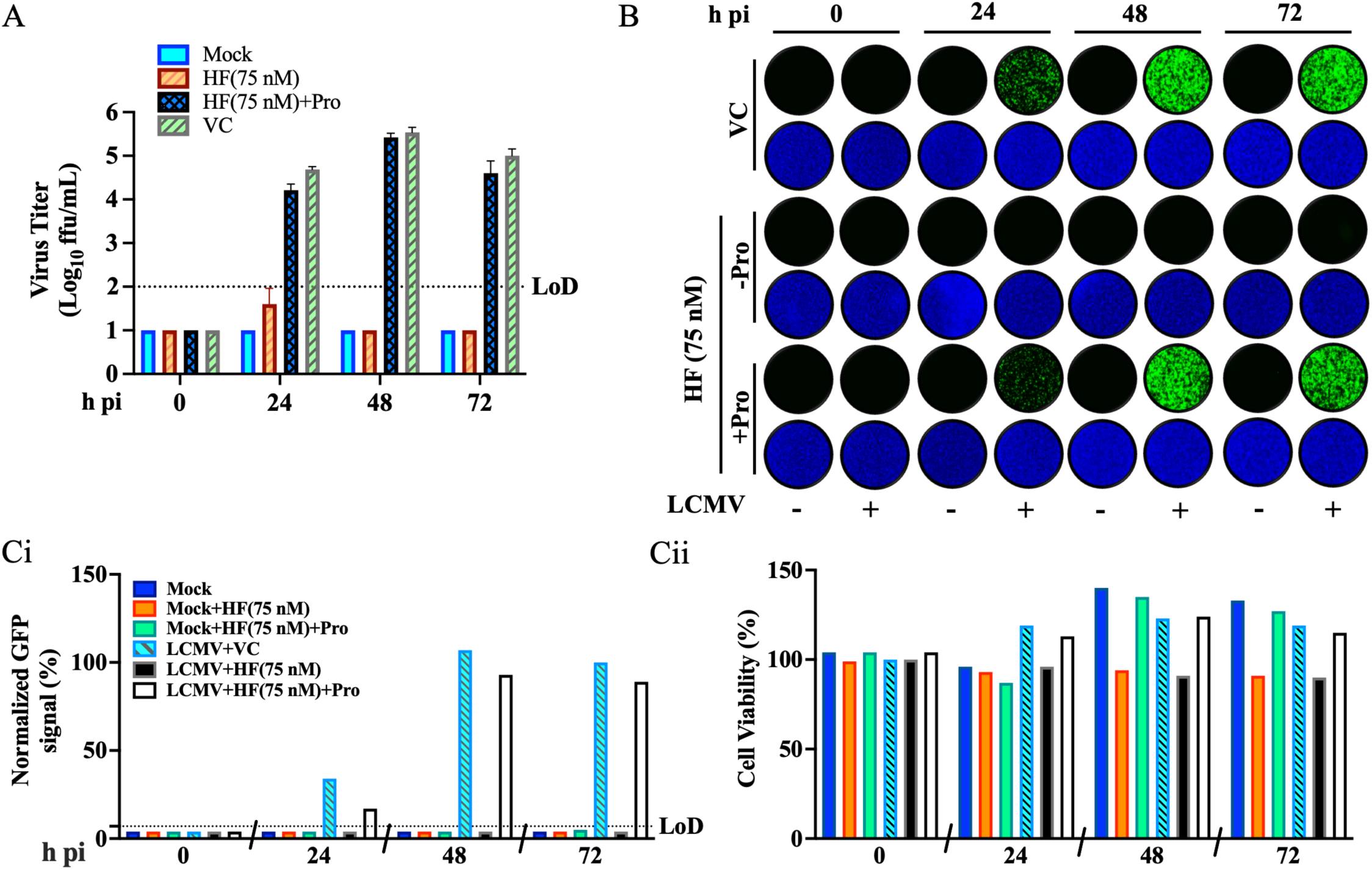
L-Proline (Pro) reversed the effect of the HF on LCMV multi-step growth kinetics in A549 cells. **A.** A549 cells were seeded at 5 × 10⁵ cells/well in an M12-well plate, infected (MOI = 0.05) with rLCMV/GFP-P2A-NP, and treated with HF (75 nM), or VC, both in the absence of presence of 1 mM Pro. At the indicated h pi, CCS were collected, and the titers of infectious virus were determined by FFA using Vero E6 cells. **B.** At the indicated h pi, samples from **A** were fixed with 4% PFA and stained with DAPI. Images (4x magnification) were collected using a Keyence BZ-X710 microscope. **C.** GFP (**Ci**) and DAPI (**Cii**) signals were quantified using Cytation 5 and normalized to the average value of the VC treated samples at 72 h pi.

### 3.4. Effect of HF on LCMV cell entry

To further investigate the mechanism by which HF exerted its antiviral activity against LCMV, we performed a time of addition experiment to assess whether HF affected a cell entry or post-cell entry step of LCMV infection. For this, we used the single-cycle infectious rLCMVΔGPC/ZsG to prevent the confounding effects of multiple rounds of replication without the need of NH4Cl treatment (Fig. 5A). Treatment with HF (25, 50 and 75 nM) at both −2h and +2h of infection significantly reduced the number of ZsG+ cells compared to VC treatment (Fig. 5A, B). In contrast, treatment with the LCMV entry inhibitor F3406 led to a reduction in ZsG⁺ cells when administered at −2h, but not at +2h, of infection. LCMV cell entry is completed in less than 60 minutes [42]. Hence, results of the time of addition assay indicated that HF treatment was able to disrupt a post-cell entry step. MaAv enter host cells through receptor-mediated endocytosis [43,44]. In the acidic environment of the endosome, GP2 facilitates a pH-dependent membrane fusion between the viral and host membranes, enabling the release of the vRNP into the cell cytoplasm, where replication and transcription occur. Therefore, the entry process cannot be completed if the GP2-mediated fusion event is inhibited. To examine whether HF was also able to interfere with GP2-mediated fusion, we transfected HEK293T with plasmids encoding LCMV GPC (pC-LCMV-GPC), LASV GPC (pC-LASV-GPC), or an empty vector (pC-E) as a control, along with a plasmid expressing GFP (pC-GFP). At 5 h post-transfection, we treated cells with HF (75 nM) or VC. At 24 h post-transfection, we exposed cells to acidic (pH 5.0) or neutral (pH 7.2) medium for 15 min and then returned them to standard medium to monitor syncytium formation by GFP fluorescence (Fig. 5C). Cells expressing LCMV or LASV GPC exhibited robust fusion activity under acidic conditions both in the absence and presence of HF, indicating that HF does not interfere with GPC-mediated pH-dependent event required to complete the virus cell entry process.

**Figure 5.**
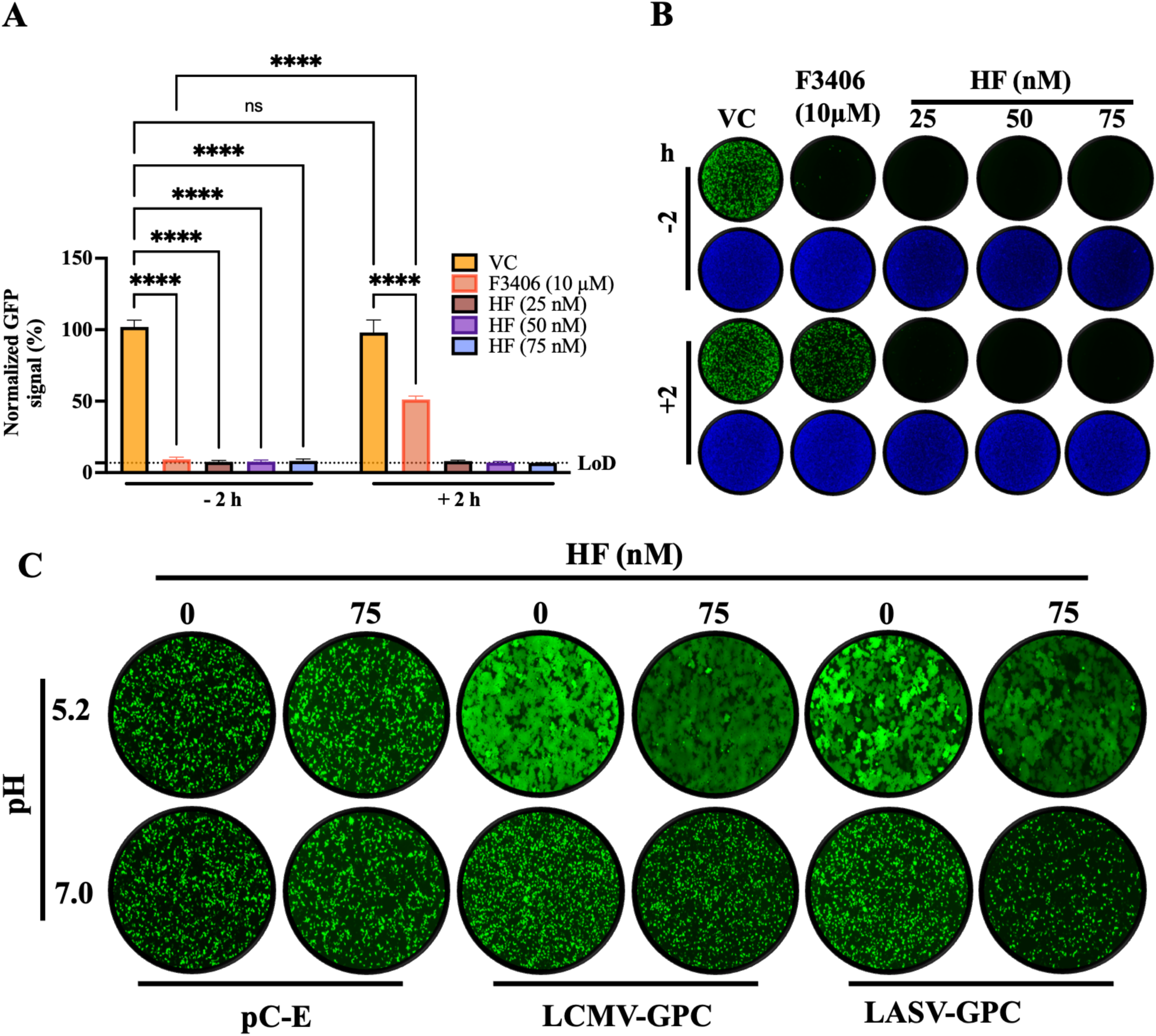
Effect of HF on LCMV cell entry. **A.** Time-of-addition assay: A549 cells were seeded in 96-well plates at 8 × 10⁴ cells/well. At 16 h post-seeding cells were infected (MOI = 0.5) with a single-cycle infectious recombinant LCMV expressing ZsGreen (rLCMVΔGPC/ZsG). HF (25, 50 or 75 nM) or VC was added either 2 h prior to infection (−2 h) or 2 h post-infection (+2 h). The LCMV entry inhibitor F3406 (10 µM) served as control. At 48 h pi, ZsGreen-positive (ZsG⁺) cells were quantified using the Cytation 5 Microplate Reader (BioTek, Agilent) and normalized (%) to VC-treated infected controls. Results correspond to the mean and SD of four biological replicates. **B.** IF images from panel **A** samples. **C.** Effect of HF on LCMV and LASV GPC-mediated fusion. HEK293T cells were seeded (8×10^5^ cells/well) onto poly-L-lysine-coated 24-well plates. At 16 h post-seeding cells were transfected (4 µg/well) with plasmids encoding GPC from LCMV (pC-LCMV-GPC), or LASV (pC-LASV-GPC), or empty pCAGGS vector (pC-E) together with a GFP-expressing plasmid (pC-GFP; 50 ng/well). At 5 h post-transfection, cells were washed with DMEM supplemented with 10% FBS and treated with either HF (75 nM) or vehicle control (VC). At 24 hours post-transfection, monolayers were exposed to acidified (pH 5.0) or neutral (pH 7.2) medium for 15 minutes and then returned to neutral medium (DMEM/10% FBS). The pattern of GFP expression was used to monitor fusion events over time, and once syncytium formation was apparent in VC-treated cells, samples were fixed (4% PFA) and IF images (4x) collected using a Keyence BZ-X710 microscope.

### 3.5. Effect of HF on LCMV vRNP activity

As with other negative strand RNA viruses, LCMV vRNP is responsible for directing the biosynthetic processes of replication and transcription of the viral genome. To evaluate the impact of HF on LCMV vRNP activity, we employed a cell-based LCMV MG system. This system recapitulates LCMV RNA replication and transcription through intracellular reconstitution of the viral vRNP, which requires co-expression of the LCMV L and NP proteins together with a plasmid expressing the MG RNA (Fig.1). Levels of MG-directed GFP expression provides an integrated readout of vRNP activity. We transfected HEK293T cells in suspension with plasmids expressing the components of a LCMV MG system and seeded them at 6×10⁴ and 8×10⁴ cells/96-well. We used a MG system where the intracellularly reconstituted vRNP directed expression of GFP. Samples where the L expressing plasmid was not included in the served as a negative control. Transfected cells were treated with HF (75 nM), Pro (1 mM), Pro (1 mM) + HF (75 nM), or VC. At 72 h post-transfection, cells were fixed with 4% PFA and stained with DAPI. Representative IF images of each treatment condition (Fig. 6A) were collected using a Keyence BZ-X710 microscope. GFP (Fig. 6Bi) and DAPI (Fig. 6Bii) signals were quantified using a Cytation 5 plate reader (BioTek, Agilent) and normalized (%) to VC-treated controls, which were set at 100% (Fig. 6Ci, Cii). Treatment with HF (75 nM) resulted in ∼50% reduction in GFP expression compared to VC-treated controls. We did not observe significant differences on the effect of HF on MG activity at the two seeding cell densities of 6 x 10^4^ and 8 x 10^4^ cells/96-well (Fig. 6D), indicating that the modest cytostatic effect associated with HF treatment did not affect the activity of LCMV vRNP.

**Figure 6.**
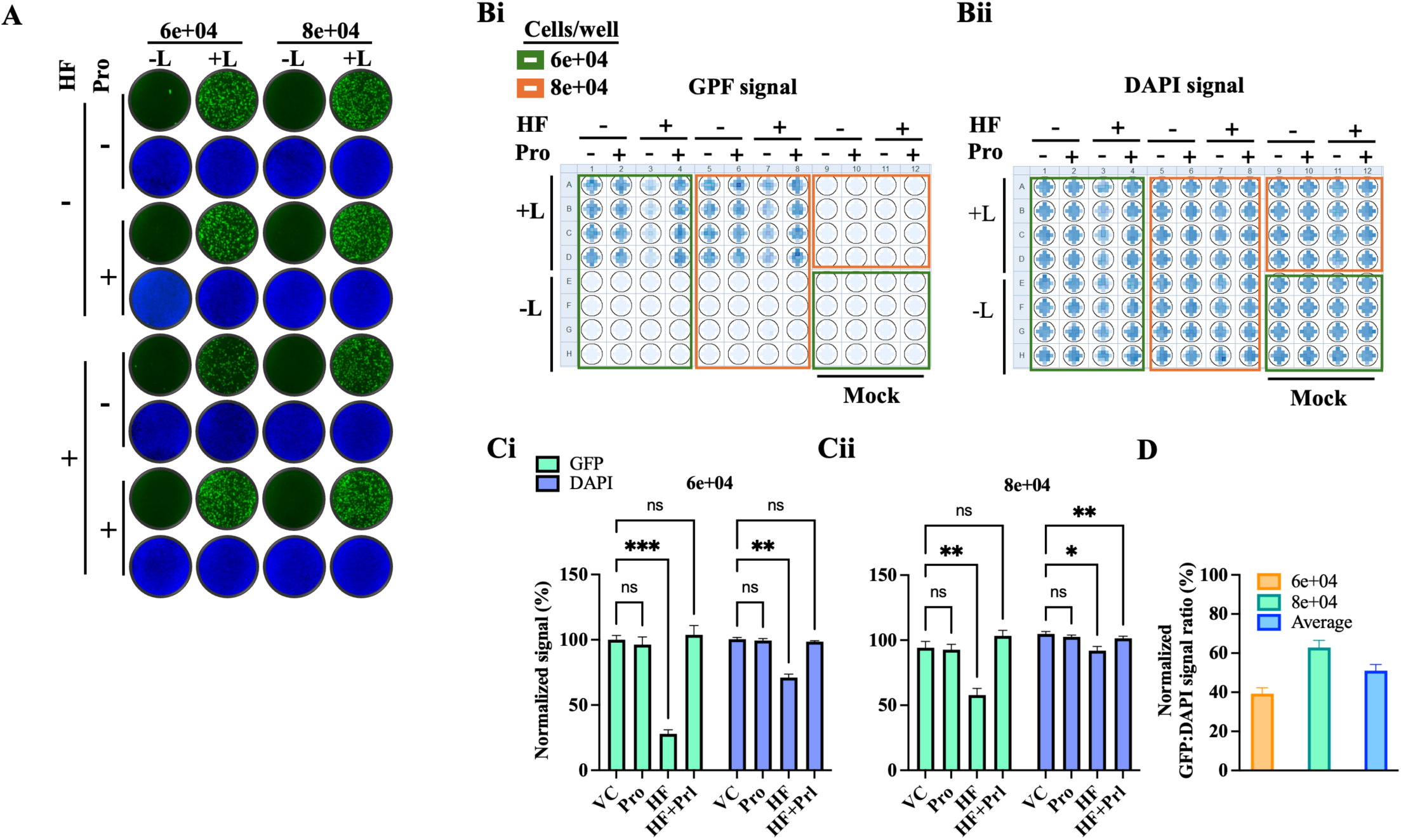
Effect of HF on LCMV vRNP activity. HEK293T cells were transfected in suspension with plasmids expressing the components of the LCMV T7-MG-GFP system (NP, L, and T7-MG-GFP). Samples where the L expressing plasmid was not included in the transfection mixture served as a negative control for the MG activity. Transfected cells were seeded (80 µL/well containing either 6×10⁴ or 8×10⁴ cells) into 96-well plates pre-spotted with 20 µL of media containing either VC, Pro (1 mM), HF (75 nM), or Pro (1 mM) + HF (75 nM), and mixed by pipetting twice to ensure homogenous distribution. At 72 h post-transfection, cells were fixed with 4% PFA and stained with DAPI. **A.** Representative IF images (4x magnification) from each treatment condition (B) were collected using a Keyence BZ-X710 microscope. **B.** GFP (**Bi)** and DAPI (**Bii**) signals were quantified using a Cytation 5 plate reader (BioTek, Agilent). C. GFP and DAPI signals were normalized (%) to VC-treated controls, which were set at 100% activity. Results represent the mean and SD of four independent replicates for 6×10⁴ (**Ci**) and 8×10⁴ (**Cii**). **D.** Normalized GFP:DAPI signal ratio of samples treated with HF (75 nM) at two different cells densities (**Ci** and **Cii**).

### 3.6. Effect of HF on LCMV assembly and budding

The MaAv matrix Z protein is the main driver of virus budding [45,46]. To determine whether HF affected Z-mediated virus budding, we used an established cell-based Z budding assay where the activity of Gluc serves as a surrogate marker for Z budding activity [47]. We transfected HEK293T cells with a plasmid encoding a chimeric GLuc where its N-terminal secretory signal peptide was replaced with the Z open reading frame (Z-GLuc) of either LCMV or LASV. Transfected cells were treated with HF (75 nM) or VC, and at 48 h post-transfection GLuc activity was measured in both CCSs, containing virus-like particles (VLPs), and whole-cell lysates (WCLs). Z budding efficiency was calculated as the ratio of VLP-associated GLuc (GLuc in CCS) to total GLuc (GLuc CCS + GLuc-WCL) x 100. Values of budding efficiency were normalized (%) to those of VC-treated cells transfected with Z-WT that were assigned a value of 100%. As control, we use Z(G2A)-GLuc, where mutation G2A prevents Z myristoylation, which results in inhibition of Z budding activity. Treatment with HF inhibited Z budding activity (Fig.7A). We next examined whether HF inhibited assembly and release of infectious particles. For this, we infected A549 cells with rLCMV/GFP-P2A-NP (MOI = 0.05) and at 24 h pi, we collected CCSs and washed cell monolayers three times with fresh media and refed them with medium containing HF (75 nM) or VC and allowed infection to proceed. At 48 h pi we collected CCSs and fixed cells with 4% PFA. Virus titers in CCSs of cells that were refed with medium containing HF were similar at 24 and 48 h pi (Fig. 7B). In contrast, in cells refed with medium without HF virus titers increased from ∼5 logs (24 h pi) to ∼6 logs (48 h pi) (Fig. 7B). This finding correlated with the restricted virus propagation between 24 and 48 h pi in infected cells that were refed with medium containing HF (Fig. 7C-E). To further investigate the effect of HF on Z protein expression and budding, we infected A549 cells with a recombinant LCMV expressing a C-terminal HA-tagged Z protein (rLCMV/Z-HA) (MOI = 1) [39] and treated them with different concentrations of HF, or with the N-myristoyltransferase inhibitor (NMTi) IMP-1088 [39] or ribavirin (RBV). At 72 h pi we determined the NP to Z ratio by IF using a mouse monoclonal anti-HA antibody to detect Z and the rat monoclonal VL4 antibody to LCMV NP. Consistent with the HF inhibitory effect on LCMV multiplication (Fig. 3 and Fig. 4), treatment with HF resulted in a dose-dependent reduction of expression levels of both Z and NP. However, the normalized NP:Z ratio decreased in HF, compared to RBV, treated infected cells, indicating that HF treatment promoted Z, but not NP, intracellular accumulation (Fig. 7F), which correlated with increased intracellular levels of GLuc (Fig 7G). As predicted, the NP:Z ratio increased in IMP-1088, compared to RBV, treated infected cells due to IMP-1088 induced proteasome mediated degradation of Z (Fig. 7Fii) [39].

**Figure 7.**
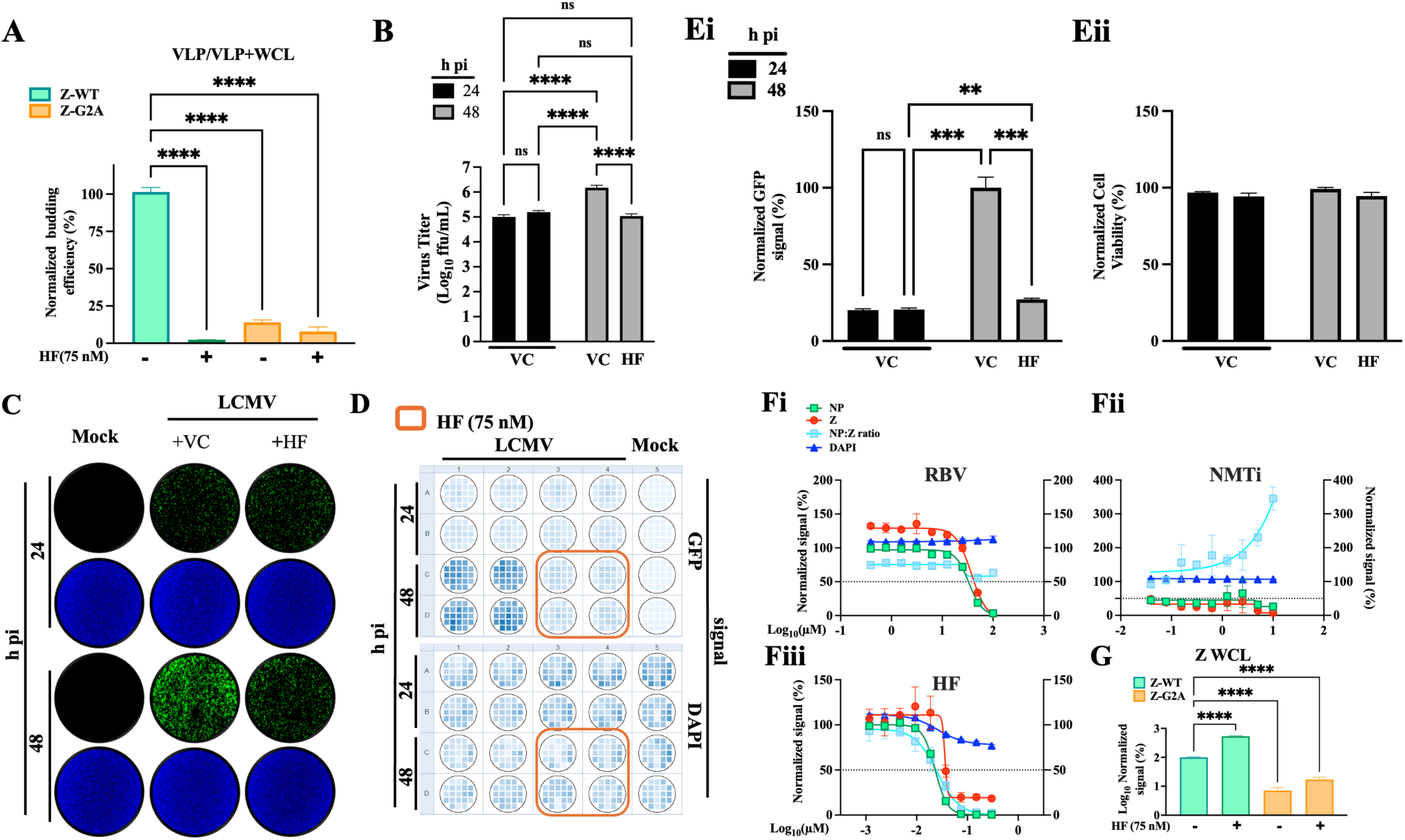
Effect of HF on LCMV Z budding activity and viral progeny release. **A.** Effect of HF on Z budding activity. HEK293T cells were seeded onto poly-L-lysine-coated 12-well plates at a density of 5 × 10⁵ cells per well. At 16 h post-seeding cells were transfected with plasmids encoding LCMV Z fused to Gaussia luciferase (GLuc) (pC-LCMV-Z-GLuc), or a myristoylation-deficient mutant form of Z (pC-LCMV-Z-G2A-GLuc). At 5 h post-transfection, cells were washed three times and incubated with fresh medium containing the indicated concentrations of the tested compounds. At 48 h post-transfection, CCS were collected and whole-cell lysates (WCL) prepared. GLuc activity in both CCS and WCL was quantified using the SteadyGlo Luciferase Pierce: Gaussia Luciferase Glow Assay Kit and measured with a Cytation 5 reader (BioTek, Agilent). GLuc activity in CCS served as a surrogate for Z-containing VLP released via Z budding activity, while GLuc activity in WCL reflected intracellular Z expression levels. Z budding efficiency was calculated as CCS-GLuc/CCS-GLuc + WCL-GLuc and normalized to VC-treated samples, set at 100%. Data were visualized using Prism 10. **B.** Effect of HF on virus progeny release. A549 cells were seeded at 8 × 10^5^ cells/well in an M12-well plate and infected (MOI = 0.05) with rLCMV/GFP-P2A-NP. At 24 h pi CCS were collected, and cells were washed 3 times with fresh media and refed with medium containing 75 nM HF (+ HF) or VC (+ VC). The same set of samples were fixed with 4% PFA and sealed to prevent the leak of PFA fumes, and after 20 minutes incubation the seal was removed, fixed cells washed 3 times with PBS and left in 2 mL/well of PBS. At 48 h pi, CCS were collected, and cells were fixed with 4% PFA. Titers of infectious virus in CCSs at 24 and 48 h pi were determined by FFA using Vero E6 cells. **C.** Cells fixed at 24 and 48 h pi were stained with DAPI. Representative IF images (4X) from each treatment condition were collected using a Keyence BZ-X710 microscope. **D.** GFP and DAPI signals were collected using Cytation 5 plate reader. **E.** GFP (**Ei**) and DAPI staining (**Eii)** signals were normalized (%) to values corresponding to 48 h pi VC-treated samples. **F.** Effect of HF treatment on Z intracellular levels. A549 cells were seeded in 96-well clear-bottom plates at a density of 8.0 × 10⁴ cells per well. Sixteen hours later, cells were infected (MOI = 1) with a recombinant LCMV expressing HA-tagged Z protein (rLCMV/Z-HA) and treated with the indicated concentrations of HF, the NMTi IMP-1088, RBV, or VC. At 72 h post-treatment, cells were fixed with 4% PFA and analyzed by IF using antibodies to HA (Z) and NP. Nuclei were visualized with DAPI staining. to quantify cell viability. Each data point represents the mean of four (HF) or two (IMP-1088 and RBV) independent replicates. **G.** GLuc activity in WCL was quantified using the SteadyGlo Luciferase Pierce: Gaussia Luciferase Glow Assay Kit and measured with a Cytation 5 reader (BioTek, Agilent).

### 3.7. Effect of HF on multiplication of LASV and JUNV in cultured cells

Related viruses often depend on shared host cell factors for replication, making these factors attractive targets for broad-spectrum antiviral strategies. To evaluate whether HF exhibited a broad-spectrum anti-MaAv activity, we tested HF’s ability to inhibit multiplication of LASV and JUNV in cell culture. For this we used tri-segmented versions of LASV (r3LASV) and JUNV (r3JUNV) expressing the GFP and GLuc reporter genes whose expression levels served as accurate readouts of virus multiplication [37,38]. HF exhibited a potent dose-dependent inhibitory effect on multiplication of LASV and JUNV in A549 cells as determined by reduced GFP and GLuc expression levels (Fig. 8).

**Figure 8.**
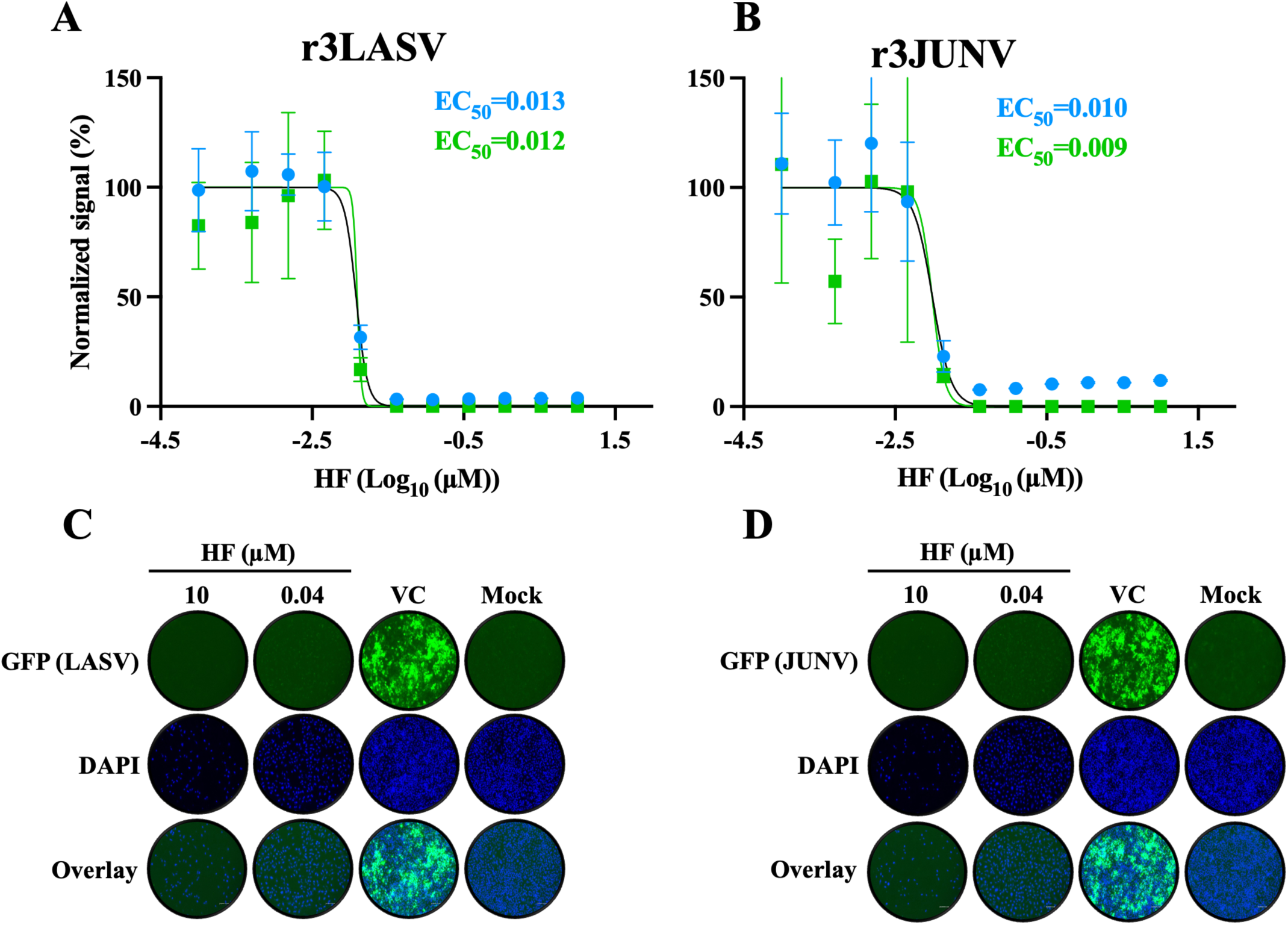
Effect of HF on multiplication of JUNV, and LASV. A549 cells were seeded in 96-well plates at a density of 2 × 10⁴ cells per well, infected (MPI = 0.05) with recombinant tri-segmented forms of LASV (r3LASV, MOI = 0.003) or JUNV (r3JUNV, MOI = 0.006) expressing GFP and GLuc and treated them with HF at the indicated concentrations. At 72 h pi CCS were collected and cells fixed with 4% PFA and subjected to DAPI staining. GFP signals and intensity of DAPI staining were determined using the Cytation 5 plate reader. Both GFP and DAPI signals were normalized (%) to VC-treated infected controls. **A, B.** EC₅₀ values based on GFP (green) and GLuc (blue) of HF for LASV (**A**) and JUNV (**B**) were calculated using a four-parameter variable slope model in Prism 10. Data represent the mean ± standard deviation (SD) of four independent biological replicates. **C, D.** Representative IF images (72 h pi) of LASV (**C**) and JUNV (**D**) infected cells and treated with the indicated concentrations of HF or VC.

MaAv share basic aspects of their molecular and cell biology, and therefore it would be expected that HF also targeted a post-cell entry step of LASV and JUNV lifecycle. To validate this prediction, we evaluated the effect of HF on the activity of LASV vRNP using a LASV cell-based MG system where MG-directed levels of GFP expression served as surrogate of the LASV vRNP activity. We transfected HEK293T cells with plasmids expressing the components of the LASV MG system and seeded them at 6×10⁴ and 8×10⁴ cells/96-well. Samples where the L expressing plasmid was not included in the transfection mixture served as a negative control. Transfected cells were treated with HF (75 nM), Pro (1 mM), Pro (1 mM) + HF (75 nM), or VC. At 72 h post-transfection, cells were fixed with 4% PFA and stained with DAPI. Representative IF images of each treatment condition (Fig. 9A) were collected using a Keyence BZ-X710 microscope. GFP (Fig. 9B) and DAPI (Fig. 9C) signals were quantified using a Cytation 5 plate reader (BioTek, Agilent) and normalized (%) to VC-treated controls, which were set at 100% (Fig. 9D, E). As with the LCMV MG assay, we observed that HF treatment caused a modest (∼40%), but significant, reduction on LASV MG activity at the two seeding cell densities of 6 x 10⁴ and 8 x 10⁴ cells/96-well, indicating that the modest cytostatic effect associated with HF treatment did not affect the activity of LASV vRNP significantly.

**Figure 9.**
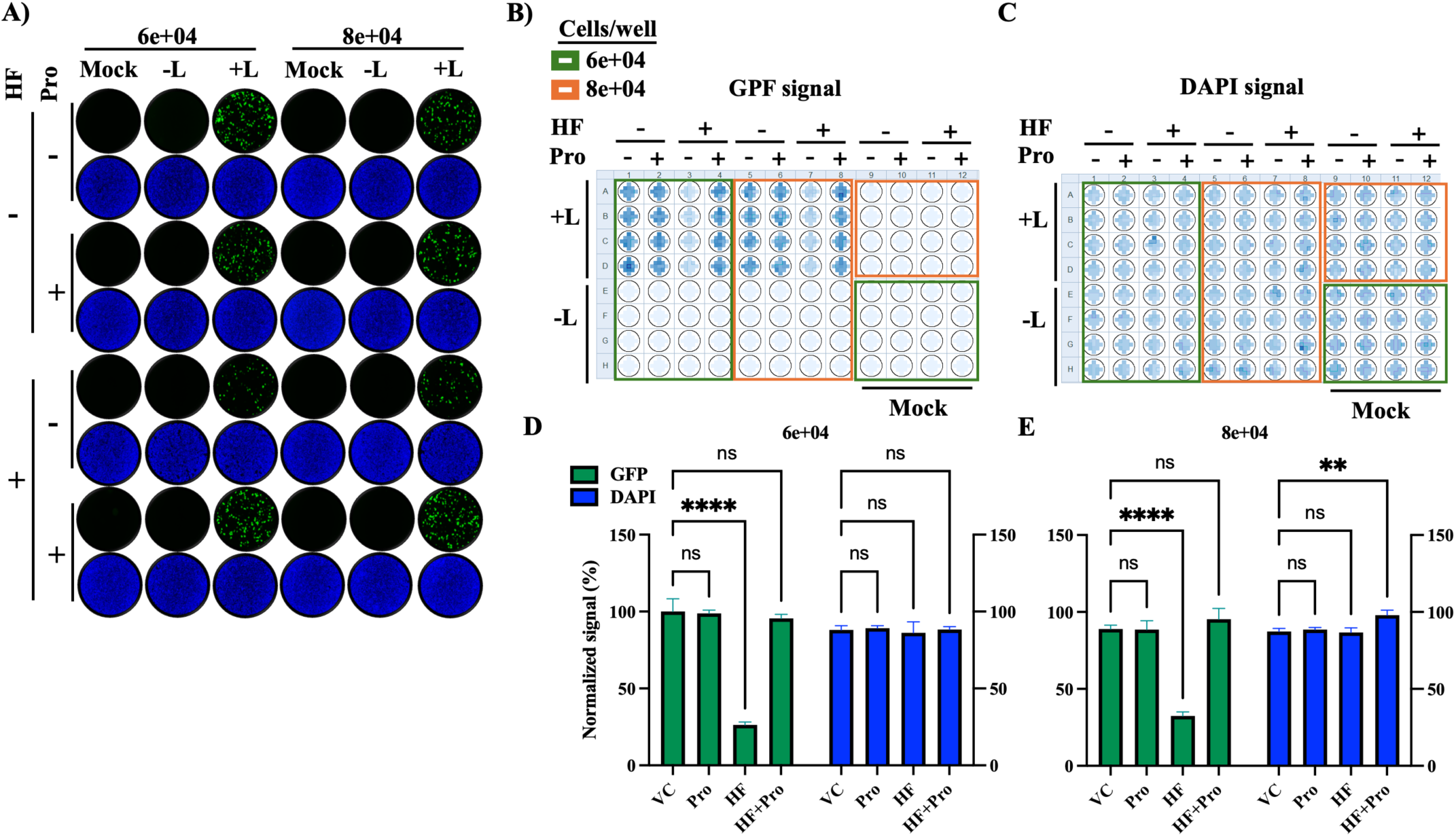
Effect of HF on LASV vRNP activity. **A.** HEK293T cells were transfected in suspension with plasmids expressing the components of the LASV T7-MG-GFP system (NP, L, and T7-MG-GFP). Samples where the L expressing plasmid was not included in the transfection mixture served as a negative control for the MG activity. Transfected cells were seeded (80 µL/well containing either 6×10⁴ or 8×10⁴ cells) into 96-well plates pre-spotted with 20 µL of media containing either VC, Pro (1 mM), HF (75 nM), or Pro+HF, and mixed by pipetting twice to ensure homogenous distribution. At 72 h post-transfection, cells were fixed with 4% PFA and stained with DAPI. GFP expression levels were normalized (%) to VC-treated controls, which were set at 100% activity. IF images (4x magnification) were collected using a Keyence BZ-X710 microscope. **B, C.** GFP (**B)** and DAPI (**C**) signals were quantified using a Cytation 5 plate reader (BioTek, Agilent). **D, E.** GFP values were normalized to VC-treated controls. Results represent the mean and SD of four independent replicates using cell densities of 6×10⁴ (**D**) and 8×10⁴ (**E**) cells/96 well.

## 4. Discussion

We have presented evidence that HF is a potent inhibitor of LCMV and the hemorrhagic fever causing MaAv LASV and JUNV. The anti-MaAv activity of HF was reversed by proline supplementation (Fig. 4), supporting that inhibition of the PRS activity of EPRS1 is a main mechanism of action by which HF exerts its anti-MaAv activity. Results from tome of addition and cell-based MG assays indicated that HF did not affect virus cell entry and had only a modest (twofold) inhibitory effect on the activity of the vRNP (Fig. 6). In contrast, HF potently inhibited Z-mediated virus assembly and budding (Fig. 7), resulting in increased intracellular levels of Z, which could mediate potent inhibition of the vRNP in infected cells, this contributing to reduced production of infectious viral progeny and restricted virus propagation. HF mediated inhibition of PRS could preferentially affect translation of proteins with proline-rich motif like the Z late (L)-domain motifs that play a critical role on Z budding activity [45,46]. Likewise, HF treatment can preferentially affect expression of several functionally proline-dependent cellular proteins that contribute to Z-mediated viral budding, including CHMP2A (ESCRT-III), CHMP4B (ESCRT-III), TSG101 (ESCRT-I), and ALIX (Accessory ESCRT protein). Specific proline residues in CHMP2A and CHMP4B mediate conformational changes necessary for ESCRT-III [48] filament assembly and membrane scission [48], whereas TSG101 [49,50] and ALIX [51] utilize proline-rich motifs for protein-protein interactions essential for ESCRT recruitment and viral budding. As with many other viruses, MaAv have evolved mechanisms to counteract the host cell innate immune responses to infection, including sensing of viral RNA by PKR [38], and induction of the type I interferon response [52]. HF mediated Inhibition of PRS leads to the accumulation of uncharged tRNAs, a biochemical event linked to the activation of the amino acid starvation (AAS) stress signal that drives autophosphorylation of the amino acid sensor GCN2 kinase, and subsequent phosphorylation of eIF2α and increased expression of ATF4, the main effector of the integrated stress response (ISR) pathway [53–58]. Evidence indicates that the cellular ISR interacts with and regulate signaling intermediates involved in the activation of innate and adaptive immune responses. Future studies, beyond the scope of the present work, will investigate whether ISR contributes to HF mediated anti-MaAv activity.

HF is already in use in veterinary medicine [59], and has shown excellent safety profile [60–67] and significant efficacy in phase I and II clinical trials for different indications including cancer [68,69], raising the possibility of its repurposing as a host-directed antiviral (HDA) against human pathogenic MaAv. HF is orally bioavailable and was shown to reach an average Cmax of ∼1.3 nM) or ∼7.4 nM after a single administration of 0.5 mg or 3.5 mg doses, respectively, in a phase I clinical trial [68]. HF long half-life (∼30 h) leads to its accumulation with two-to three-fold higher exposure by day 15 of dosing [70]. In mice HF was widely distributed in tissues and its exposure in lung was > 87-fold compared to plasma after a single intravenous injection in mice [71]. This suggests that although it may be difficult to reach HF plasma levels > EC50 values we determined in this study, doses tested in phase I trials may be sufficient to reach HF therapeutic levels in tissues. HF also was shown to reduce acute inflammatory events in acute myocarditis and acute hepatitis in viral infection [72,73], which could also provide benefits against hemorrhagic fever causing MaAv infections.

HDAs represent a promising yet underexplored approach in contemporary antiviral research [74,75]. To date, most HDAs are focused on modulating interferon signaling or targeting cellular receptors, but the systematic exploration of broad-spectrum HDAs remains largely unexplored. Unlike direct-acting antivirals (DAAs) that target viral proteins and functions, HDAs disrupt host factors and cellular processes that viruses hijack for their multiplication and pathogenesis. Since members of a virus family share host dependencies, HDAs have the potential to act as broad-spectrum antivirals. In addition, HDAs pose a higher genetic barrier to the emergence of drug-resistant viral variants, which often compromise antiviral therapy [76,77]. HDAs have been shown to effectively inhibit viral infection induced by immunosuppression associated with JAK inhibitors drugs [78]. HDAs, like HF, can facilitate drug repurposing strategies for drugs with established safety profiles, thus reducing longer development timelines and regulatory barriers associated with the development of novel drugs [79]. HDA-based strategies face the challenge of risks associated with toxicity and unintended consequences of disruption of cell physiological processes. However, therapies against acute viral infections, such as hemorrhagic fever diseases caused by MaAv, would require short-term treatment, which increases the likelihood of identifying effective therapeutic regimens with acceptable safety profiles. Moreover, transient reduced levels of AARS appear to have minimal effects in global translation, and heterozygous carriers for recessive inactivating AARS mutations linked to hypomyelinating leukodystrophy do not display disease phenotypes [80]. In addition to HF, other PRS inhibitors are being evaluated in humans, including DWN12088 [81] that is in phase II clinical trials in the US and South Korea [82], and anticipated to be used for the treatment of interstitial pulmonary fibrosis. Thus, novel PRS inhibitors, including HF analogs, may lead to the identification of additional broad-spectrum antiviral agents that could be potentially repurposed as HDAs against human pathogenic MaAv.

## Author Contributions

Conceptualization, H.W. and J.C.d.l.T.; methodology, H.W., P.I., R.Z., N.J., R.E., B.C., R.S.; software, H.W.; validation, H.W., P.I., R.Z., N.J., R.E.; formal analysis, H.W., J.C.d.l.T.; investigation, H.W.; resources, L.M-S., J.C.d.l.T.; data curation, NA; writing—original draft preparation, H.W., R.K., J.C.d.l.T.; writing—review and editing, H.W., P.I., R.Z., N.J., R.E., B.C., R.S., L.M-S., J.C.d.l.T.; visualization, H.W.; supervision, H.W. and J.C.d.l.T.; project administration, H.W.; funding acquisition, L.M-S., J.C.d.l.T. All authors have read and agreed to the published version of the manuscript.

## Institutional Review Board Statement

NA.

## Informed Consent Statement

NA.

## Data Availability Statement

Data included in the article.

---

**SF1.**
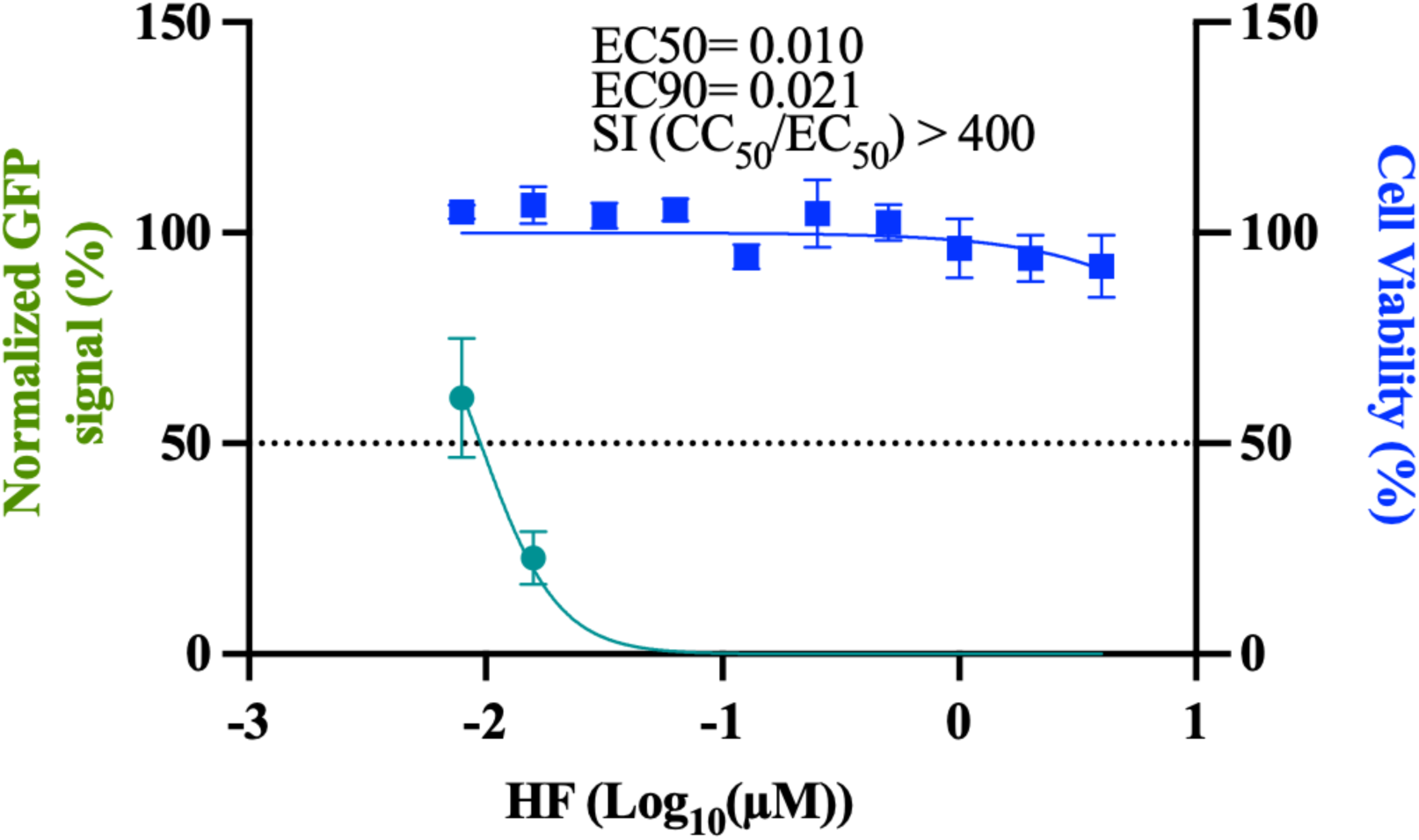
Dose-dependent effect of HF on LCMV multiplication in endothelial cells. Ea.hy926 cells were seeded at 4 × 10⁴ cells/well into a 96-well plate, infected (MOI = 0.05) with rLCMV/WT-GFP-P2A-NP and treated with HF at the indicated concentrations. At 48 h pi, cells were fixed with 4% PFA, and numbers of infected cells determined by IF. Numbers of infected cells were normalized (%) to those of VC-treated samples cells and expressed as a % of infected cells (A). Cell viability was estimated based on DAPI staining signal quantified using the Cytation 5 reader. Results correspond to the average of four biological replicates. EC50 and CC50 values were calculated using a variable slope (based on four parameters) model and EC90 values were calculated using FindECanything model (logEC50=logECF - (1/HillSlope)*log(F/(10-F)) with F parameter set to 10 (Prism10). Results show the mean and SD of four biological replicates.

**SF2.**
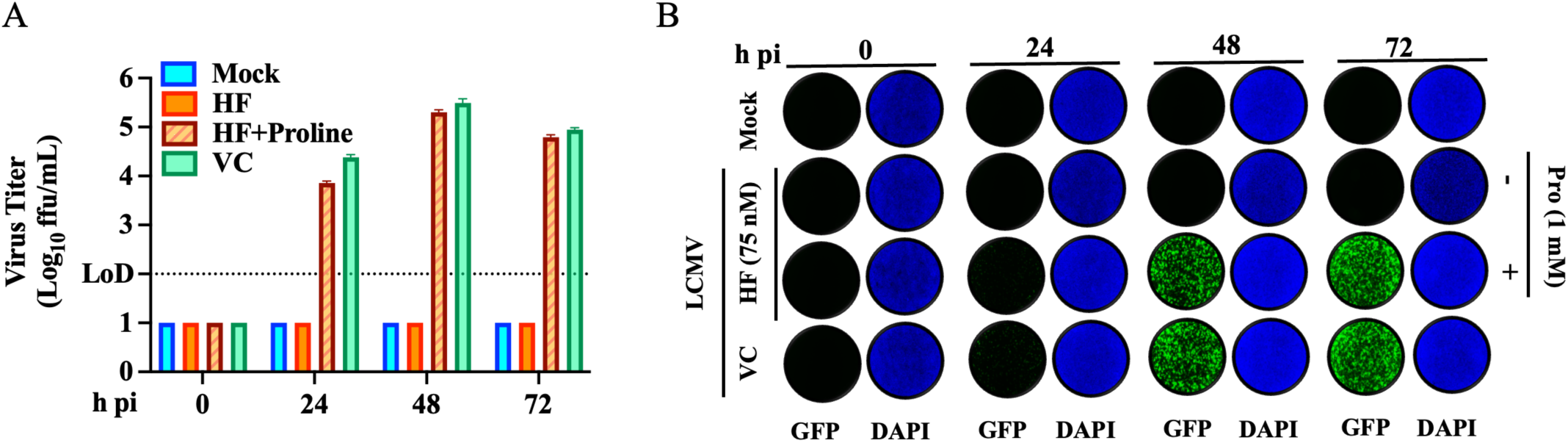
Biological replicate of HF (75 nM) on LCMV multi-step growth kinetics and peak titers in A549 cells. **A.** The counter effect of L-proline (Pro) on the production of infectious viral progeny in presence of HF. A549 cells were seeded at 5 × 10⁵ cells/well in an M12-well plate, infected with rLCMV/GFP-P2A-NP (MOI 0.05), and treated with HF (75 nM), Pro (1 mM), the combination of HF and Pro, or with VC. At the indicated time points, cell culture supernatants were collected, and the titers of infectious virus were determined by the focus-forming assay (FFA) using Vero E6 cells. **B.** At the indicated h pi, samples from A were fixed with 4%PFA and washed with DPBS, sealed and stained with DAPI at the end of the experiment and imaged at 4x magnification using Keyence BZ-X710 series.

**SF3.**
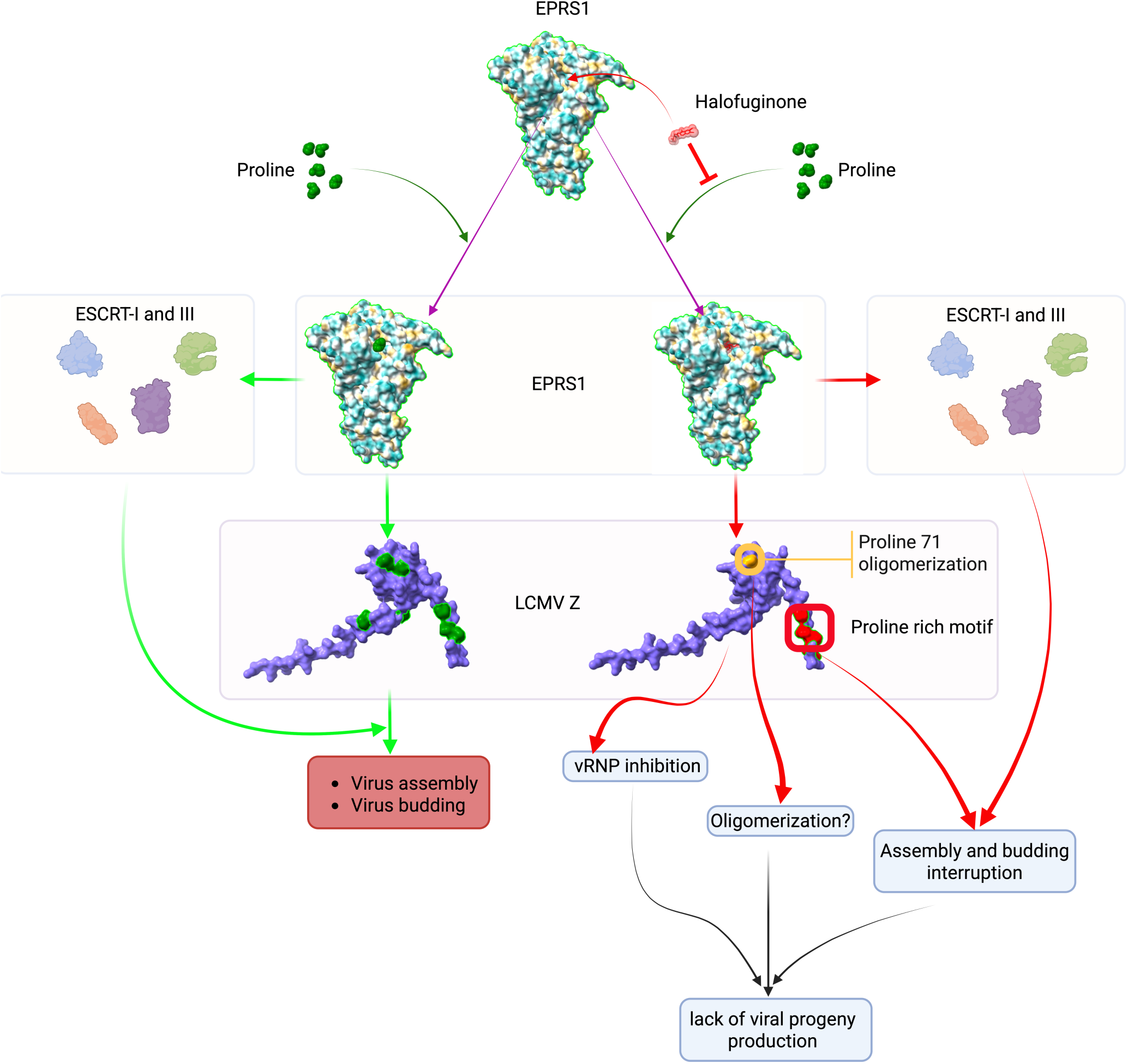
Proposed model of the effect of the prolyl tRNA synthetase 1 (PRS1) inhibitor, halofuginone (HF), on mammarenavirus assembly and budding. The PRS1 domain of the bifunctional enzyme glutamyl-prolyl-tRNA synthetase (EPRS1) catalyzes the incorporation of proline into both cellular and viral proteins. The mammarenavirus Z matrix protein orchestrates the viral assembly and budding processes. Proline-riche late domains critically contribute to Z budding activity via their interaction with components of cellular ESCRT machinery. Inhibition of PRS1 by HF disrupts Z-mediated viral assembly and budding processes, thereby blocking virus multiplication. HF-mediated inhibition of PRS1 inhibition also interferes with the activity of ESCRT-I and III components, which further contributes to disrupt Z-mediated budding. EPRS1 is depicted based on the PDB structure 4k88 [83]; LCMV Z matrix protein is presented as predicted using AlphaFold 3 server [84]. Red colored residues indicate the proline-rich motif late domains of Z matrix protein; orange residue corresponds to proline at position 71 implicated in Z oligomerization. The 3D figure was generated using ChimeraX v1.10.1. The protein sizes’ are not for scale.

## References

1. Basinski, A.J.; Fichet-Calvet, E.; Sjodin, A.R.; Varrelman, T.J.; Remien, C.H.; Layman, N.C.; Bird, B.H.; Wolking, D.J.; Monagin, C.; Ghersi, B.M.;, et al. Bridging the Gap: Using Reservoir Ecology and Human Serosurveys to Estimate Lassa Virus Spillover in West Africa. PLOS Computational Biology 2021, 17, e1008811, doi:10.1371/journal.pcbi.1008811.

2. Fichet-Calvet, E.; Rogers, D.J. Risk Maps of Lassa Fever in West Africa. PLOS Neglected Tropical Diseases 2009, 3, e388, doi:10.1371/journal.pntd.0000388.

3. Grant, D.S.; Samuels, R.J.; Garry, R.F.; Schieffelin, J.S. Lassa Fever Natural History and Clinical Management. Curr Top Microbiol Immunol 2023, 440, 165–192, doi:10.1007/82_2023_263.

4. Radoshitzky, S.R.; Buchmeier, M.; de la Torre, J.C. Emerging Viruses: Arenaviridae. In Fields Virology; Knipe, D. avid, Howley, P., Whelan, S., Eds.; 2020; Vol. I ISBN 978-1-9751-1254-7.

5. Sogoba, N.; Feldmann, H.; Safronetz, D. Lassa Fever in West Africa: Evidence for an Expanded Region of Endemicity. Zoonoses Public Health 2012, 59 *Suppl 2*, 43–47, doi:10.1111/j.1863-2378.2012.01469.x.

6. Freedman, D.O.; Woodall, J. Emerging Infectious Diseases and Risk to the Traveler. Med Clin North Am 1999, 83, 865–883, v.

7. Isaäcson, M. Viral Hemorrhagic Fever Hazards for Travelers in Africa. Clin Infect Dis 2001, 33, 1707–1712, doi:10.1086/322620.

8. Grange, Z.L.; Goldstein, T.; Johnson, C.K.; Anthony, S.; Gilardi, K.; Daszak, P.; Olival, K.J.; O’Rourke, T.; Murray, S.; Olson, S.H.;, et al. Ranking the Risk of Animal-to-Human Spillover for Newly Discovered Viruses. Proc Natl Acad Sci U S A 2021, 118, e2002324118, doi:10.1073/pnas.2002324118.

9. Grant, A.; Seregin, A.; Huang, C.; Kolokoltsova, O.; Brasier, A.; Peters, C.; Paessler, S. Junín Virus Pathogenesis and Virus Replication. Viruses 2012, 4, 2317–2339, doi:10.3390/v4102317.

10. Lendino, A.; Castellanos, A.A.; Pigott, D.M.; Han, B.A. A Review of Emerging Health Threats from Zoonotic New World Mammarenaviruses. BMC Microbiology 2024, 24, 115, doi:10.1186/s12866-024-03257-w.

11. Borio, L.; Inglesby, T.; Peters, C.J.; Schmaljohn, A.L.; Hughes, J.M.; Jahrling, P.B.; Ksiazek, T.; Johnson, K.M.; Meyerhoff, A.; O’Toole, T.;, et al. Hemorrhagic Fever Viruses as Biological Weapons: Medical and Public Health Management. JAMA 2002, 287, 2391–2405, doi:10.1001/jama.287.18.2391.

12. Bonthius, D.J. Lymphocytic Choriomeningitis Virus: A Prenatal and Postnatal Threat. Adv Pediatr 2009, 56, 75–86, doi:10.1016/j.yapd.2009.08.007.

13. Palacios, G.; Druce, J.; Du, L.; Tran, T.; Birch, C.; Briese, T.; Conlan, S.; Quan, P.-L.; Hui, J.; Marshall, J.;, et al. A New Arenavirus in a Cluster of Fatal Transplant-Associated Diseases. N Engl J Med 2008, 358, 991–998, doi:10.1056/nejmoa073785.

14. Pencole, L.; Sibiude, J.; Weingertner, A.S.; Mandelbrot, L.; Vauloup-Fellous, C.; Picone, O. Congenital Lymphocytic Choriomeningitis Virus: A Review. Prenat Diagn 2022, 42, 1059–1069, doi:10.1002/pd.6192.

15. Lapošová, K.; Pastoreková, S.; Tomášková, J. Lymphocytic Choriomeningitis Virus: Invisible but Not Innocent. Acta Virol 2013, 57, 160–170, doi:10.4149/av_2013_02_160.

16. Gass, J.T.; Nofchissey, R.A.; Twohig, F.M.; Ye, C.; Goodfellow, S.M.; Mentore, K.; Burgos, M.; Negrete, O.; Whitmer, S.; Klena, J.D.;, et al. Discovery of a Novel Lymphocytic Choriomeningitis Virus Strain Associated with Severe Human Disease in Immunocompetent Patient, New Mexico. Emerg Microbes Infect 2025, 14, 2542250, doi:10.1080/22221751.2025.2542250.

17. Salam, A.P.; Cheng, V.; Edwards, T.; Olliaro, P.; Sterne, J.; Horby, P. Time to Reconsider the Role of Ribavirin in Lassa Fever. PLOS Neglected Tropical Diseases 2021, 15, e0009522, doi:10.1371/journal.pntd.0009522.

18. Cross, R.W.; Hastie, K.M.; Mire, C.E.; Robinson, J.E.; Geisbert, T.W.; Branco, L.M.; Ollmann Saphire, E.; Garry, R.F. Antibody Therapy for Lassa Fever. Curr Opin Virol 2019, 37, 97–104, doi:10.1016/j.coviro.2019.07.003.

19. Gowen, B.B.; Juelich, T.L.; Sefing, E.J.; Brasel, T.; Smith, J.K.; Zhang, L.; Tigabu, B.; Hill, T.E.; Yun, T.; Pietzsch, C.;, et al. Favipiravir (T-705) Inhibits Junín Virus Infection and Reduces Mortality in a Guinea Pig Model of Argentine Hemorrhagic Fever. PLOS Neglected Tropical Diseases 2013, 7, e2614, doi:10.1371/journal.pntd.0002614.

20. Mendenhall, M.; Russell, A.; Juelich, T.; Messina, E.L.; Smee, D.F.; Freiberg, A.N.; Holbrook, M.R.; Furuta, Y.; de la Torre, J.-C.; Nunberg, J.H.;, et al. T-705 (Favipiravir) Inhibition of Arenavirus Replication in Cell Culture. Antimicrob Agents Chemother 2011, 55, 782–787, doi:10.1128/AAC.01219-10.

21. Rosenke, K.; Feldmann, H.; Westover, J.B.; Hanley, P.W.; Martellaro, C.; Feldmann, F.; Saturday, G.; Lovaglio, J.; Scott, D.P.; Furuta, Y.;, et al. Use of Favipiravir to Treat Lassa Virus Infection in Macaques. Emerg Infect Dis 2018, 24, 1696–1699, doi:10.3201/eid2409.180233.

22. Safronetz, D.; Rosenke, K.; Westover, J.B.; Martellaro, C.; Okumura, A.; Furuta, Y.; Geisbert, J.; Saturday, G.; Komeno, T.; Geisbert, T.W.;, et al. The Broad-Spectrum Antiviral Favipiravir Protects Guinea Pigs from Lethal Lassa Virus Infection Post-Disease Onset. Sci Rep 2015, 5, 14775, doi:10.1038/srep14775.

23. Lieber, C.M.; Plemper, R.K. 4’-Fluorouridine Is a Broad-Spectrum Orally Available First-Line Antiviral That May Improve Pandemic Preparedness. DNA Cell Biol 2022, 41, 699–704, doi:10.1089/dna.2022.0312.

24. Welch, S.R.; Spengler, J.R.; Westover, J.B.; Bailey, K.W.; Davies, K.A.; Aida-Ficken, V.; Bluemling, G.R.; Boardman, K.M.; Wasson, S.R.; Mao, S.;, et al. Delayed Low-Dose Oral Administration of 4′-Fluorouridine Inhibits Pathogenic Arenaviruses in Animal Models of Lethal Disease. Science Translational Medicine 2024, 16, eado7034, doi:10.1126/scitranslmed.ado7034.

25. Cashman, K.A.; Wilkinson, E.R.; Posakony, J.; Madu, I.G.; Tarcha, E.J.; Lustig, K.H.; Korth, M.J.; Bedard, K.M.; Amberg, S.M. Lassa Antiviral LHF-535 Protects Guinea Pigs from Lethal Challenge. Sci Rep 2022, 12, 19911, doi:10.1038/s41598-022-23760-2.

26. Gowen, B.B.; Naik, S.; Westover, J.B.; Brown, E.R.; Gantla, V.R.; Fetsko, A.; Dagley, A.L.; Blotter, D.J.; Anderson, N.; McCormack, K.;, et al. Potent Inhibition of Arenavirus Infection by a Novel Fusion Inhibitor. Antiviral Res 2021, 193, 105125, doi:10.1016/j.antiviral.2021.105125.

27. C, E.; Oo, A.; A, M.; K, O.; O, E.; E, E.; Tg, A.; N, A.; C, A.; S, O.; et al. Electrocardiographic Alterations in Patients Treated for Acute Lassa Fever: Description of Results from a Phase II Clinical Trial in Nigeria. Journal of Infection and Public Health 2025, 18.

28. Andersen, K.G.; Shapiro, B.J.; Matranga, C.B.; Sealfon, R.; Lin, A.E.; Moses, L.M.; Folarin, O.A.; Goba, A.; Odia, I.; Ehiane, P.E.;, et al. Clinical Sequencing Uncovers Origins and Evolution of Lassa Virus. Cell 2015, 162, 738–750, doi:10.1016/j.cell.2015.07.020.

29. Zhang, G.; Cao, J.; Cai, Y.; Liu, Y.; Li, Y.; Wang, P.; Guo, J.; Jia, X.; Zhang, M.; Xiao, G.;, et al. Structure-Activity Relationship Optimization for Lassa Virus Fusion Inhibitors Targeting the Transmembrane Domain of GP2. Protein Cell 2019, 10, 137–142, doi:10.1007/s13238-018-0604-x.

30. de Jonge, M.J.A.; Dumez, H.; Verweij, J.; Yarkoni, S.; Snyder, D.; Lacombe, D.; Marréaud, S.; Yamaguchi, T.; Punt, C.J.A.; van Oosterom, A.;, et al. Phase I and Pharmacokinetic Study of Halofuginone, an Oral Quinazolinone Derivative in Patients with Advanced Solid Tumours. Eur J Cancer 2006, 42, 1768–1774, doi:10.1016/j.ejca.2005.12.027.

31. Koon, H.B.; Fingleton, B.; Lee, J.Y.; Geyer, J.T.; Cesarman, E.; Parise, R.A.; Egorin, M.J.; Dezube, B.J.; Aboulafia, D.; Krown, S.E. Phase II AIDS Malignancy Consortium Trial of Topical Halofuginone in AIDS-Related Kaposi Sarcoma. J Acquir Immune Defic Syndr 2011, 56, 64–68, doi:10.1097/QAI.0b013e3181fc0141.

32. Pines, M.; Snyder, D.; Yarkoni, S.; Nagler, A. Halofuginone to Treat Fibrosis in Chronic Graft-versus-Host Disease and Scleroderma. Biol Blood Marrow Transplant 2003, 9, 417–425, doi:10.1016/s1083-8791(03)00151-4.

33. García-Rodríguez, I.; Moreni, G.; Capendale, P.E.; Mulder, L.; Aknouch, I.; Vieira de Sá, R.; Johannesson, N.; Freeze, E.; van Eijk, H.; Koen, G.;, et al. Assessment of the Broad-Spectrum Host Targeting Antiviral Efficacy of Halofuginone Hydrobromide in Human Airway, Intestinal and Brain Organotypic Models. Antiviral Res 2024, 222, 105798, doi:10.1016/j.antiviral.2024.105798.

34. Hwang, J.; Jiang, A.; Fikrig, E. A Potent Prolyl tRNA Synthetase Inhibitor Antagonizes Chikungunya and Dengue Viruses. Antiviral Res 2019, 161, 163–168, doi:10.1016/j.antiviral.2018.11.017.

35. Wu, J.; Subbaiah, K.C.V.; Xie, L.H.; Jiang, F.; Khor, E.-S.; Mickelsen, D.; Myers, J.R.; Tang, W.H.W.; Yao, P. Glutamyl-Prolyl-tRNA Synthetase Regulates Proline-Rich Pro-Fibrotic Protein Synthesis During Cardiac Fibrosis. Circulation research 2020, 127, 827, doi:10.1161/CIRCRESAHA.119.315999.

36. Iwasaki, M.; Minder, P.; Caì, Y.; Kuhn, J.H.; Yates, J.R.; Torbett, B.E.; de la Torre, J.C. Interactome Analysis of the Lymphocytic Choriomeningitis Virus Nucleoprotein in Infected Cells Reveals ATPase Na+/K+ Transporting Subunit Alpha 1 and Prohibitin as Host-Cell Factors Involved in the Life Cycle of Mammarenaviruses. PLoS Pathog 2018, 14, e1006892, doi:10.1371/journal.ppat.1006892.

37. Emonet, S.F.; Seregin, A.V.; Yun, N.E.; Poussard, A.L.; Walker, A.G.; de la Torre, J.C.; Paessler, S. Rescue from Cloned cDNAs and In Vivo Characterization of Recombinant Pathogenic Romero and Live-Attenuated Candid #1 Strains of Junin Virus, the Causative Agent of Argentine Hemorrhagic Fever Disease. Journal of Virology 2011, 85, 1473–1483, doi:10.1128/jvi.02102-10.

38. Witwit, H.; Khafaji, R.; Salaniwal, A.; Kim, A.S.; Cubitt, B.; Jackson, N.; Ye, C.; Weiss, S.R.; Martinez-Sobrido, L.; de la Torre, J.C. Activation of Protein Kinase Receptor (PKR) Plays a pro-Viral Role in Mammarenavirus-Infected Cells. J Virol 2024, 98, e0188323, doi:10.1128/jvi.01883-23.

39. Witwit, H.; Betancourt, C.A.; Cubitt, B.; Khafaji, R.; Kowalski, H.; Jackson, N.; Ye, C.; Martinez-Sobrido, L.; de la Torre, J.C. Cellular N-Myristoyl Transferases Are Required for Mammarenavirus Multiplication. Viruses 2024, 16, 1362, doi:10.3390/v16091362.

40. Battegay, M.; Cooper, S.; Althage, A.; Bänziger, J.; Hengartner, H.; Zinkernagel, R.M. Quantification of Lymphocytic Choriomeningitis Virus with an Immunological Focus Assay in 24-or 96-Well Plates. J Virol Methods 1991, 33, 191–198, doi:10.1016/0166-0934(91)90018-u.

41. Perez, M.; de la Torre, J.C. Characterization of the Genomic Promoter of the Prototypic Arenavirus Lymphocytic Choriomeningitis Virus. Journal of Virology 2003, 77, 1184–1194, doi:10.1128/jvi.77.2.1184-1194.2003.

42. Quirin, K.; Eschli, B.; Scheu, I.; Poort, L.; Kartenbeck, J.; Helenius, A. Lymphocytic Choriomeningitis Virus Uses a Novel Endocytic Pathway for Infectious Entry via Late Endosomes. Virology 2008, 378, 21–33, doi:10.1016/j.virol.2008.04.046.

43. Borrow, P.; Oldstone, M.B. Mechanism of Lymphocytic Choriomeningitis Virus Entry into Cells. Virology 1994, 198, 1–9, doi:10.1006/viro.1994.1001.

44. White, J.; Helenius, A. pH-Dependent Fusion between the Semliki Forest Virus Membrane and Liposomes. Proc Natl Acad Sci U S A 1980, 77, 3273–3277, doi:10.1073/pnas.77.6.3273.

45. Perez, M.; Greenwald, D.L.; de La Torre, J.C. Myristoylation of the RING Finger Z Protein Is Essential for Arenavirus Budding. Journal of Virology 2004, 78, 11443–11448, doi:10.1128/jvi.78.20.11443-11448.2004.

46. Witwit, H.; de la Torre, J.C. Mammarenavirus Z Protein Myristoylation and Oligomerization Are Not Required for Its Dose-Dependent Inhibitory Effect on vRNP Activity. BioChem 2025, 5, 10, doi:10.3390/biochem5020010.

47. Capul, A.A.; de la Torre, J.C. A Cell-Based Luciferase Assay Amenable to High-Throughput Screening of Inhibitors of Arenavirus Budding. Virology 2008, 382, 107–114, doi:10.1016/j.virol.2008.09.008.

48. Scott, A.; Gaspar, J.; Stuchell-Brereton, M.D.; Alam, S.L.; Skalicky, J.J.; Sundquist, W.I. Structure and ESCRT-III Protein Interactions of the MIT Domain of Human VPS4A. Proceedings of the National Academy of Sciences 2005, 102, 13813–13818, doi:10.1073/pnas.0502165102.

49. Garrus, J.E.; Schwedler, U.K. von; Pornillos, O.W.; Morham, S.G.; Zavitz, K.H.; Wang, H.E.; Wettstein, D.A.; Stray, K.M.; Côté, M.; Rich, R.L.;, et al. Tsg101 and the Vacuolar Protein Sorting Pathway Are Essential for HIV-1 Budding. Cell 2001, 107, 55–65, doi:10.1016/S0092-8674(01)00506-2.

50. Martin-Serrano, J.; Zang, T.; Bieniasz, P.D. HIV-1 and Ebola Virus Encode Small Peptide Motifs That Recruit Tsg101 to Sites of Particle Assembly to Facilitate Egress. Nat Med 2001, 7, 1313–1319, doi:10.1038/nm1201-1313.

51. Strack, B.; Calistri, A.; Craig, S.; Popova, E.; Göttlinger, H.G. AIP1/ALIX Is a Binding Partner for HIV-1 P6 and EIAV P9 Functioning in Virus Budding. Cell 2003, 114, 689–699, doi:10.1016/S0092-8674(03)00653-6.

52. Martínez-Sobrido, L.; Emonet, S.; Giannakas, P.; Cubitt, B.; García-Sastre, A.; de la Torre, J.C. Identification of Amino Acid Residues Critical for the Anti-Interferon Activity of the Nucleoprotein of the Prototypic Arenavirus Lymphocytic Choriomeningitis Virus. J Virol 2009, 83, 11330–11340, doi:10.1128/JVI.00763-09.

53. Rai, S.; Szaruga, M.; Pitera, A.P.; Bertolotti, A. Integrated Stress Response Activator Halofuginone Protects Mice from Diabetes-like Phenotypes. J Cell Biol 2024, 223, e202405175, doi:10.1083/jcb.202405175.

54. Pitera, A.P.; Szaruga, M.; Peak-Chew, S.; Wingett, S.W.; Bertolotti, A. Cellular Responses to Halofuginone Reveal a Vulnerability of the GCN2 Branch of the Integrated Stress Response. The EMBO Journal 2022, 41, e109985, doi:10.15252/embj.2021109985.

55. Flores, M.E.; McNamara-Bordewick, N.K.; Lovinger, N.L.; Snow, J.W. Halofuginone Triggers a Transcriptional Program Centered on Ribosome Biogenesis and Function in Honey Bees. Insect Biochemistry and Molecular Biology 2021, 139, 103667, doi:10.1016/j.ibmb.2021.103667.

56. Kamberov, Y.G.; Kim, J.; Mazitschek, R.; Kuo, W.P.; Whitman, M. Microarray Profiling Reveals the Integrated Stress Response Is Activated by Halofuginone in Mammary Epithelial Cells. BMC Res Notes 2011, 4, 381, doi:10.1186/1756-0500-4-381.

57. Kates, M.; St. Paul, M.; Ohashi, P.S.; Saibil, S.D. Protocol for Halofuginone-Mediated Metabolic Reprogramming of Murine T Cells via Activation of the GCN2 Pathway. STAR Protocols 2025, 6, 104173, doi:10.1016/j.xpro.2025.104173.

58. Proto-Siqueira, R.; Santos, M.G.; Carvalho, V.M.; Maekawa, Y.H.; Testagrossa, L.A.; Andrade, Z.R.; Oliveira, J.S.; Chauffaille, M. de L.L.F.; Zago, M.A.; Nagler, A.;, et al. Halofuginone Induces Post-Transcriptional Down-Regulation of Cyclin D1, Cell Cycle Arrest and Apoptosis In Mantle Cell Lymphoma Cells through Activation of Integrated Stress Response Pathways. Blood 2010, 116, 773, doi:10.1182/blood.V116.21.773.773.

59. Daugschies, A.; Gässlein, U.; Rommel, M. Comparative Efficacy of Anticoccidials under the Conditions of Commercial Broiler Production and in Battery Trials. Veterinary Parasitology 1998, 76, 163–171, doi:10.1016/S0304-4017(97)00203-3.

60. Zhang, D.-F.; Sun, B.-B.; Yue, Y.-Y.; Yu, H.-J.; Zhang, H.-L.; Zhou, Q.-J.; Du, A.-F. Anticoccidial Effect of Halofuginone Hydrobromide against Eimeria Tenella with Associated Histology. Parasitol Res 2012, 111, 695–701, doi:10.1007/s00436-012-2889-7.

61. De Waele, V.; Speybroeck, N.; Berkvens, D.; Mulcahy, G.; Murphy, T.M. Control of Cryptosporidiosis in Neonatal Calves: Use of Halofuginone Lactate in Two Different Calf Rearing Systems. Prev Vet Med 2010, 96, 143–151, doi:10.1016/j.prevetmed.2010.06.017.

62. Crosley, R.I.; Casey, N.H.; Smith, G.A.; Roosendaal, B. Influence of Phased Inclusion of Halofuginone on Broiler Skin Tensile Strength and Growth Performance. J S Afr Vet Assoc 1992, 63, 11–15.

63. Leiba, M.; Jakubikova, J.; Klippel, S.; Mitsiades, C.S.; Hideshima, T.; Tai, Y.-T.; Leiba, A.; Pines, M.; Richardson, P.G.; Nagler, A.;, et al. Halofuginone Inhibits Multiple Myeloma Growth in Vitro and in Vivo and Enhances Cytotoxicity of Conventional and Novel Agents. Br J Haematol 2012, 157, 718–731, doi:10.1111/j.1365-2141.2012.09120.x.

64. Jarvie, B.D.; Trotz-Williams, L.A.; McKnight, D.R.; Leslie, K.E.; Wallace, M.M.; Todd, C.G.; Sharpe, P.H.; Peregrine, A.S. Effect of Halofuginone Lactate on the Occurrence of Cryptosporidium Parvum and Growth of Neonatal Dairy Calves. J Dairy Sci 2005, 88, 1801–1806, doi:10.3168/jds.S0022-0302(05)72854-X.

65. Giadinis, N.D.; Papadopoulos, E.; Panousis, N.; Papazahariadou, M.; Lafi, S.Q.; Karatzias, H. Effect of Halofuginone Lactate on Treatment and Prevention of Lamb Cryptosporidiosis: An Extensive Field Trial. J Vet Pharmacol Ther 2007, 30, 578–582, doi:10.1111/j.1365-2885.2007.00900.x.

66. Lefay, D.; Naciri, M.; Poirier, P.; Chermette, R. Efficacy of Halofuginone Lactate in the Prevention of Cryptosporidiosis in Suckling Calves. Vet Rec 2001, 148, 108–112, doi:10.1136/vr.148.4.108.

67. Almawly, J.; Prattley, D.; French, N.P.; Lopez-Villalobos, N.; Hedgespeth, B.; Grinberg, A. Utility of Halofuginone Lactate for the Prevention of Natural Cryptosporidiosis of Calves, in the Presence of Co-Infection with Rotavirus and Salmonella Typhimurium. Vet Parasitol 2013, 197, 59–67, doi:10.1016/j.vetpar.2013.04.029.

68. de Jonge, M.J.A.; Dumez, H.; Verweij, J.; Yarkoni, S.; Snyder, D.; Lacombe, D.; Marréaud, S.; Yamaguchi, T.; Punt, C.J.A.; van Oosterom, A. Phase I and Pharmacokinetic Study of Halofuginone, an Oral Quinazolinone Derivative in Patients with Advanced Solid Tumours. European Journal of Cancer 2006, 42, 1768–1774, doi:10.1016/j.ejca.2005.12.027.

69. Koon, H.B.; Fingleton, B.; Lee, J.Y.; Geyer, J.T.; Cesarman, E.; Parise, R.A.; Egorin, M.J.; Dezube, B.J.; Aboulafia, D.; Krown, S.E. Phase II AIDS Malignancy Consortium Trial of Topical Halofuginone in AIDS-Related Kaposi Sarcoma. JAIDS Journal of Acquired Immune Deficiency Syndromes 2011, 56, 64, doi:10.1097/QAI.0b013e3181fc0141.

70. de Jonge, M.J.A.; Dumez, H.; Verweij, J.; Yarkoni, S.; Snyder, D.; Lacombe, D.; Marréaud, S.; Yamaguchi, T.; Punt, C.J.A.; van Oosterom, A.;, et al. Phase I and Pharmacokinetic Study of Halofuginone, an Oral Quinazolinone Derivative in Patients with Advanced Solid Tumours. Eur J Cancer 2006, 42, 1768–1774, doi:10.1016/j.ejca.2005.12.027.

71. Stecklair, K.P.; Hamburger, D.R.; Egorin, M.J.; Parise, R.A.; Covey, J.M.; Eiseman, J.L. Pharmacokinetics and Tissue Distribution of Halofuginone (NSC 713205) in CD2F1 Mice and Fischer 344 Rats. Cancer Chemother Pharmacol 2001, 48, 375–382, doi:10.1007/s002800100367.

72. Sun, X.-H.; Fu, J.; Sun, D.-Q. Halofuginone Alleviates Acute Viral Myocarditis in Suckling BALB/c Mice by Inhibiting TGF-Β1. Biochemical and Biophysical Research Communications 2016, 473, 558–564, doi:10.1016/j.bbrc.2016.03.118.

73. Zhan, W.; Kang, Y.; Chen, N.; Mao, C.; Kang, Y.; Shang, J. Halofuginone Ameliorates Inflammation in Severe Acute Hepatitis B Virus (HBV)-Infected SD Rats through AMPK Activation. Drug Des Devel Ther 2017, 11, 2947–2955, doi:10.2147/DDDT.S149623.

74. Kumar, N.; Sharma, S.; Kumar, R.; Tripathi, B.N.; Barua, S.; Ly, H.; Rouse, B.T. Host-Directed Antiviral Therapy. Clin Microbiol Rev 2020, 33, e00168–19, doi:10.1128/CMR.00168-19.

75. Zumla, A.; Rao, M.; Wallis, R.S.; Kaufmann, S.H.E.; Rustomjee, R.; Mwaba, P.; Vilaplana, C.; Yeboah-Manu, D.; Chakaya, J.; Ippolito, G.;, et al. Host-Directed Therapies for Infectious Diseases: Current Status, Recent Progress, and Future Prospects. Lancet Infect Dis 2016, 16, e47–63, doi:10.1016/S1473-3099(16)00078-5.

76. Domingo, E.; Martin, V.; Perales, C.; Grande-Pérez, A.; García-Arriaza, J.; Arias, A. Viruses as Quasispecies: Biological Implications. Curr Top Microbiol Immunol 2006, 299, 51–82, doi:10.1007/3-540-26397-7_3.

77. Kai, Y.; Hikita, H.; Morishita, N.; Murai, K.; Nakabori, T.; Iio, S.; Hagiwara, H.; Imai, Y.; Tamura, S.; Tsutsui, S.;, et al. Baseline Quasispecies Selection and Novel Mutations Contribute to Emerging Resistance-Associated Substitutions in Hepatitis C Virus after Direct-Acting Antiviral Treatment. Sci Rep 2017, 7, 41660, doi:10.1038/srep41660.

78. Witwit, H.; de la Torre, J.C. N-Myristoyltransferase Inhibitors as Candidate Broad-Spectrum Antivirals to Treat Viral Infections Promoted by Immunosuppression Associated with JAK Inhibitors Therapy. Antiviral Research 2025, 242, 106258, doi:10.1016/j.antiviral.2025.106258.

79. Pushpakom, S.; Iorio, F.; Eyers, P.A.; Escott, K.J.; Hopper, S.; Wells, A.; Doig, A.; Guilliams, T.; Latimer, J.; McNamee, C.;, et al. Drug Repurposing: Progress, Challenges and Recommendations. Nat Rev Drug Discov 2019, 18, 41–58, doi:10.1038/nrd.2018.168.

80. Kwon, N.H.; Fox, P.L.; Kim, S. Aminoacyl-tRNA Synthetases as Therapeutic Targets. Nat Rev Drug Discov 2019, 18, 629–650, doi:10.1038/s41573-019-0026-3.

81. Yoon, I.; Kim, S.; Cho, M.; You, K.A.; Son, J.; Lee, C.; Suh, J.H.; Bae, D.; Kim, J.M.; Oh, S.;, et al. Control of Fibrosis with Enhanced Safety via Asymmetric Inhibition of prolyl-tRNA Synthetase 1. EMBO Mol Med 2023, 15, EMMM202216940, doi:10.15252/emmm.202216940.

82. Daewoong Pharmaceutical Co. LTD. A Phase 2, Randomized, Double-Blind, Placebo-Controlled Study to Evaluate the Safety and Efficacy of DWN12088 in Patients With Idiopathic Pulmonary Fibrosis; clinicaltrials.gov, 2025;

83. Son, J.; Lee, E.H.; Park, M.; Kim, J.H.; Kim, J.; Kim, S.; Jeon, Y.H.; Hwang, K.Y. Conformational Changes in Human Prolyl-tRNA Synthetase upon Binding of the Substrates Proline and ATP and the Inhibitor Halofuginone. Acta Cryst D 2013, 69, 2136–2145, doi:10.1107/S0907444913020556.

84. Abramson, J.; Adler, J.; Dunger, J.; Evans, R.; Green, T.; Pritzel, A.; Ronneberger, O.; Willmore, L.; Ballard, A.J.; Bambrick, J.;, et al. Accurate Structure Prediction of Biomolecular Interactions with AlphaFold 3. Nature 2024, 630, 493–500, doi:10.1038/s41586-024-07487-w.

